# Controlling reversible cell differentiation for labor division in microbial consortia

**DOI:** 10.1101/2021.08.03.454926

**Authors:** Davide Salzano, Davide Fiore, Mario di Bernardo

## Abstract

We address the problem of regulating and keeping at a desired balance the relative numbers between cells exhibiting a different phenotype within a monostrain microbial consortium. We propose a strategy based on the use of external control inputs, assuming each cell in the community is endowed with a reversible, bistable memory mechanism. Specifically, we provide a general analytical framework to guide the design of external feedback control strategies aimed at balancing the ratio between cells whose memory is stabilized at either one of two equilibria associated to different cell phenotypes. We demonstrate the stability and robustness properties of the control laws proposed and validate them in silico by means of a realistic agent-based implementation enabling cooperative bioproduction in microbial consortia. The proposed control framework may be used to allow long term coexistence of different populations, with both industrial and biotechnological applications.

## 1 Introduction

Synthetic Biology aims at engineering biological systems with new functionalities [1], with applications ranging from health treatments to bioremediation [2], production of biofuels and drugs in bioreactors [3]. This is made possible by embedding artificial genetic circuits into living cells, such as bacteria, yeast, and fungi, modifying their natural behavior [4]; that is, by synthetically modifying when and how much genes are expressed to produce proteins or other chemicals of interest. However, the level of complexity and the functions of such engineered genetic circuits are limited by intrinsic factors in the host cells, such as possible excessive metabolic burden, competition of limited resources and incompatible chemical reactions [5]. To overcome these limitations, a promising strategy is to distribute the required functionalities among multiple cell populations forming a microbial consortium, so that each cell strain embeds a smaller subset of engineered gene networks. [6–9]. In this way, each cell population carries out a specialized function and, by dividing labor with the others in the consortium, contributes more efficiently to the achievement of the overall final goal.

Unfortunately, this solution introduces an additional challenge; cells expressing different genes might also grow and divide at different rates. In particular, cells in the consortium whose function is associated to a lower metabolic burden will grow faster, eventually becoming dominant over the other populations; thus compromising the overall function of the consortium and giving rise to undesired dynamics, such as oscillations, or even extinction [10]. Therefore, it is crucial to develop methods to guarantee the stable coexistence between cell populations in a consortium by regulating and maintaining their relative numbers to some desired level, adjusting it to the requirements of the specific application of interest. This is possible, as suggested in [11], by using *feedback control algorithms* able to sense the relative size of all the populations involved and respond by applying appropriate stimuli to the cells in order to regulate their relative numbers. We proposed to term this problem as *ratiometric control* of cell populations [11] as its overall goal is to achieve and maintain a certain desired ratio between the size of the populations in the consortium, despite differences in their growth rates, noise and perturbations (Figure 1.c). Examples of external stimuli that can be applied to this aim include changes in the concentration of some inducer molecules in the growth medium or light stimuli applied via optogenetics.

**Figure 1:**
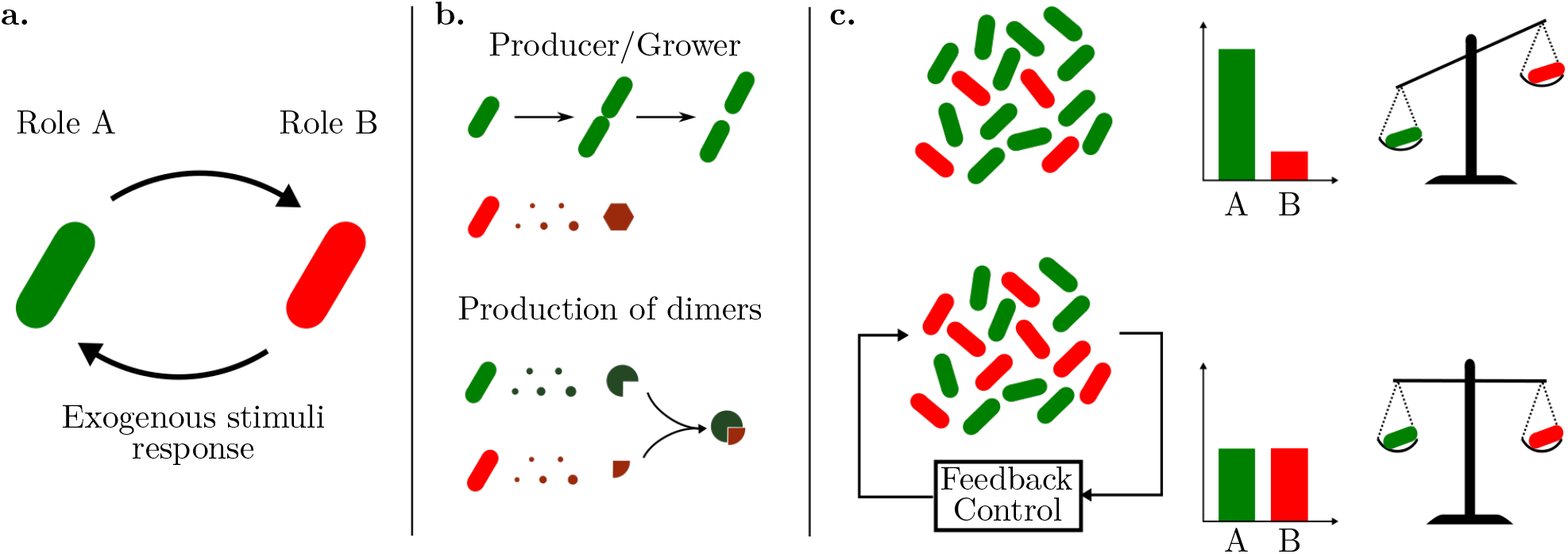
Microbial consortia composed of reversible differentiable cells can be balanced in real-time by means of external feedback controllers to guarantee efficient labor division. (**a**) Reversible differentiable cells can carry out different roles by activating/deactivating specific set of genes, depending on which state of the internal bistable memory is currently active. Cells can change role in response to exogenous stimuli from the environment, e.g., injection of inducer molecules or light. (**b**) Cells can, for example, either grow and duplicate or produce some desired molecule, or they can produce two different molecules that react and produce the desired final bioproduct. (**c**) Cells expressing different genes grow also at different rates, and thus their coexistence can be compromised. Feedback control algorithms can be employed to regulate in real-time the relative number of cells in the two groups, so that a balance in the population numbers and in the expression of desired genes is always guaranteed.

Different solutions to regulate the relative size of coexisting cell populations have been proposed in the literature, mostly based on embedding additional genetic circuitry in the cells that make them able to sense and respond to each other relative size [12–17]. Specifically, by sensing the density of the other group, cells can either increase their growth rate by producing some growth regulator protein (e.g., as in Ecolibrium project, iGEM 2016) or decrease their number by means of toxin-antitoxin mechanisms (e.g., as in [12]). Unfortunately, these embedded solutions cannot avoid the possible extinction of one of the two species, which causes either uncontrolled growth of the survivor species or its death, and are not flexible because the desired steady-state ratio of the two cell populations is hard-coded into the gene regulatory networks designed *ad hoc* and cannot be changed online. Moreover, in industrial applications where high efficiency is required, external control strategies [18–20] could be preferred to more sophisticated embedded solutions, because additional synthetic genes in the cells can cause lower production rates of the desired chemicals due to excessive metabolic load on the cells.

In this paper, to solve this problem, we consider a microbial consortium composed by *reversible differentiable cells* [21], that is, cells that belong to the same strain and embed a genetic mechanism allowing them to keep memory of past states and adapt their behavior to external stimuli from the environment, for example by activating/deactivating specific set of genes. Specifically, we consider here the simplest case of cells that can switch between two states, mimicking a flip-flop or binary memory element (Figure 1.a). The state of this bistable memory encodes the current role played by the cells in the consortium, and therefore the set of genes they are expressing at that moment. For example, a cell can use its resources either to produce some molecule or to grow and divide sustaining the cell population number [9] (Figure 1.b, top panel). Also, in the case of a genetic pathway divided into two parts, a cell can switch from one state to the other so as to activate either two depending on the overall production levels in the consortium (Figure 1.b, bottom panel). We find that by endowing the cell population with a reversible bistable system, an external control strategy can be used to solve the ratiometric control problem. Specifically, by applying external stimuli to all the cells in the consortium it is possible to switch some cells from one state to the other so as to maintain the desired ratio. We show that this is possible in a number of different ways (namely, by using relay and PI controllers) and provide stability analysis of the resulting closed-loop system and exhaustive *in silico* validation of its performance and robustness. The validation is conducted by means of agent-based simulations in BSim [22, 23], a powerful platform for realistic *in silico* experiments in bacterial populations. As a representative example we consider the realistic agent-based implementation of the proposed ratiometric control strategy to enable cooperative bioproduction in microbial consortia, showing its effectiveness and flexibility when cell growth, cell-to-cell variability and other effects are appropriately modelled.

## 2 Results

### 2.1 Reversible differentiable cells can be controlled to a desired state via a common exogenous input

The cells we consider here are assumed to embed some mechanism that can store the memory of past events. In particular, we suppose they can be switched between two different states by appropriate external stimuli, so that their *bistable* behavior can be captured at the macroscopic level by the nonlinear dynamical system (see STAR Methods 7.1 for further details)

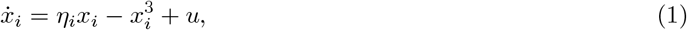

where 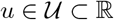 is an input signal and is in *common* to all cells.

In equation (1), 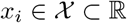 represents the macroscopic state, or role, of cell *i* and the value of parameter *η*_*i*_ ∈ ℝ is assumed to be different for all cells, accounting for their heterogeneous responses to the common external input signal *u*. For positive values of *η*_*i*_, the equation 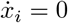 with *u* = 0 has two stable, non-trivial solutions, one negative and another positive, that we denote as *A*_*i*_ and *B*_*i*_, respectively (Figure 2.a). These solutions are the stable equilibrium points of the dynamical system described in (1) when no input is applied. We define as 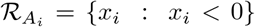 and 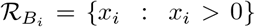 the regions of attraction [24, Sec. 8.2] of *A*_*i*_ and *B*_*i*_, respectively. Each cell will asymptotically converge to either *A*_*i*_ or *B*_*i*_ depending on which region of attraction its initial condition belongs to. Moreover, we denote by 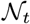 the finite set of all cells in the consortium at time *t* and with 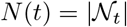 its cardinality, that is, the number of cells currently under observation (e.g., via a fluorescence microscope). Note that this number may vary in time as a consequence of cell growth or death, or because of their removal (e.g., flush away) from the culture chamber in which they are hosted. We define the sets 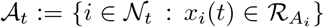 and 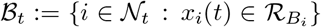, such that 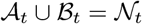 and 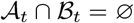, and denote with *n*_*A*_(*t*) and *n*_*B*_(*t*) their cardinality. These two sets represent the group of cells in the consortium that, at time instant *t*, in the absence of any control input *u* are expected to asymptotically converge to *A*_*i*_ and *B*_*i*_, respectively. Note that, as 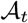 and 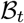 form a partition of 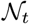, at any time it holds that *n*_*A*_(*t*) + *n*_*B*_(*t*) = *N*(*t*).

**Figure 2:**
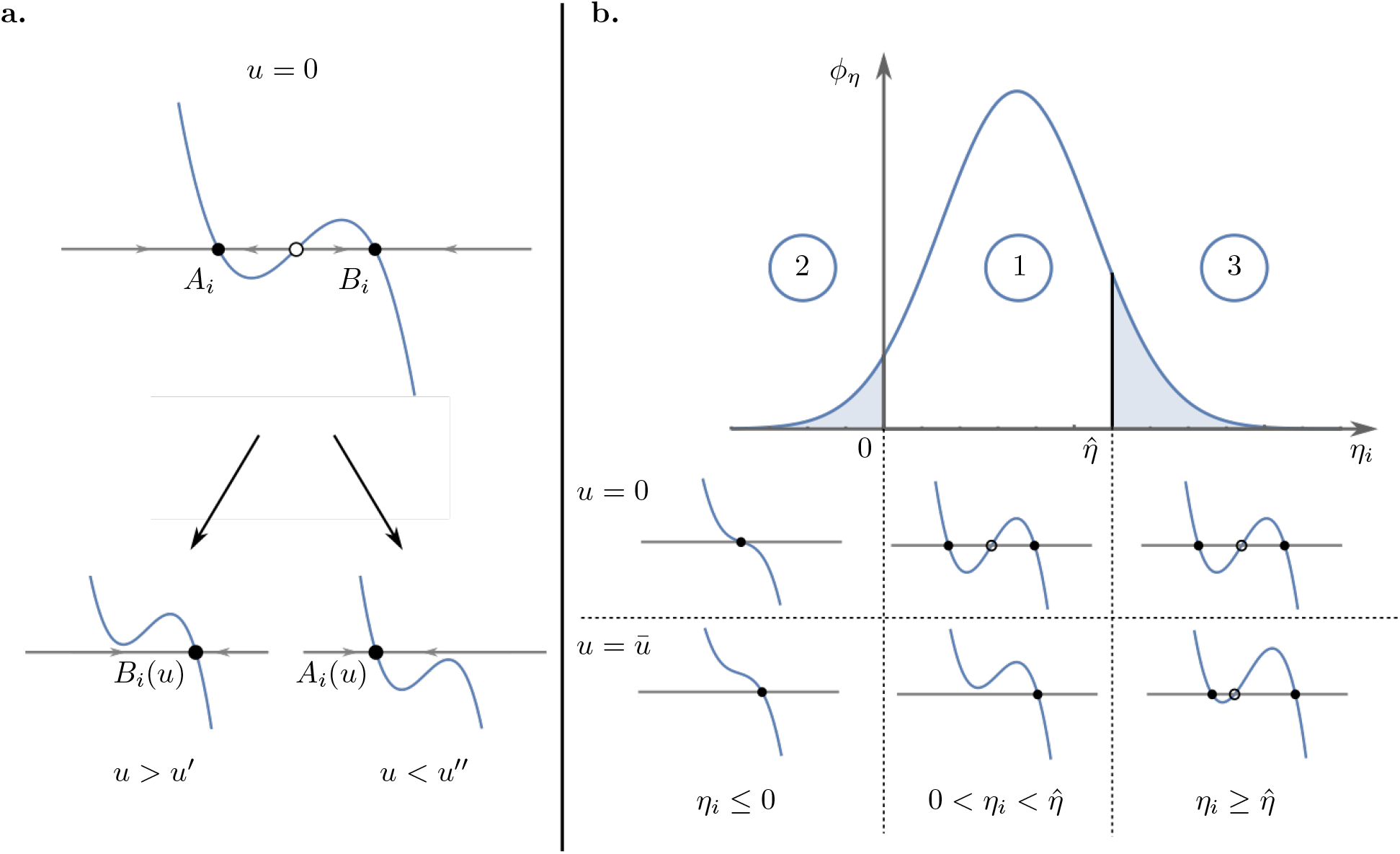
Reversible cells can be switched from state *A* to *B*, and vice versa, by means of a common external input. (**a**) The scalar dynamical system 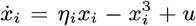, with *η*_*i*_ > 0, has two stable equilibrium points for *u* = 0, namely *A*_*i*_ and *B*_*i*_, each one corresponding to one of the two possible *roles* the cell can play in the consortium. A cell is *controllable* if by varying the input *u* in its interval of definition 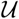, it can be moved from one group to the other and vice versa. That is, there exists an admissible value 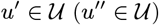 such that there is a unique positive (negative) stable solution to the equation 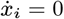 when *u* > *u′*(*u < u′′*). Full and empty dots represent stable and saddle equilibria, respectively. (**b**) Not all cells might respond as desired due to their heterogeneity, captured here by different values of the parameter *η*_*i*_ (assumed to be drawn from some probability distribution with density function *ϕ*_*η*_, here sketched as Gaussian, just for the sake of illustration). Only cells whose value of the parameter *η*_*i*_ is between 0 and 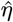 are *controllable* (Case 1), that is, they have two stable equilibria for *u* = 0 and a unique stable equilibrium for *u* = ±*ū*. Others cells can either be memory-less, or *monostable*, that is, they have only one equilibrium point for all values of *u* (Case 2), or can be *unswitchable*, having two stable equilibria for every 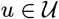, and therefore cannot change role in the consortium (Case 3).

We model cell-to-cell variability by assuming that the parameter *η*_*i*_ in (1) is drawn randomly for each cell from the real interval 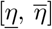 with some probability distribution (Figure 2.b). Also, we assume that the control input *u* is upper bounded by some maximum value 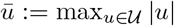, corresponding, for example, to the highest concentration of the exogenous input that can be provided by the experimental platform. In the presence of such a bound on the control signal, only cells whose parameter value *η*_*i*_ is smaller than the threshold value 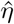 defined as

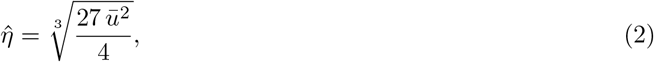

can be switched from one state to the other by an admissible value of *u*. We define cells fulfilling the condition 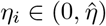 as *controllable* cells.

All the cells not fulfilling this condition are instead uncontrollable because they are either *monostable* – their parameters differing so much from their nominal values that their bistable nature is lost – or *unswitchable* because their parameters exceed the threshold value (2). In the former case *η*_*i*_ will be taking non-positive values in our model, that is, *η*_*i*_ ≤ 0, while in the latter case 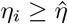.

Therefore all cells characterized by parameter values 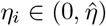, can be switched from one state to the other by means of a common bounded external input *u* applied into the environment. It is therefore possible to design, for such subset of controllable cells, some feedback control law to automatically regulate their state and keep the balance in the consortium between the two groups 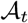 and 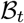 to some desired level. As we are going to show next, uncontrollable cells will contribute to a small residual error that can be precisely estimated as a function of the upper bound *ū* on the control input and therefore appropriately taken into account in the applications.

### 2.2 Ratiometric control of cell populations can be achieved by using external feedback strategies

The goal of ratiometric control is to regulate and maintain the *relative ratios* between the number of cells in 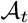 and the number of cells in 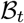, defined as

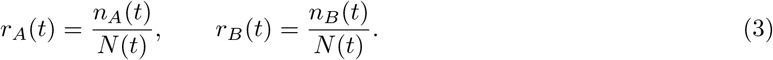

As by definition, *r*_*A*_(*t*) + *r*_*B*_(*t*) = 1 for all time, it suffices to control either *r*_*A*_ or *r*_*B*_ to control the other. Without loss of generality, we assume the ratio *r*(*t*) to be controlled is *r*_*B*_(*t*).

More formally, the objective of the *ratiometric control problem* can be stated as follows.

#### Objective

*Given a consortium of reversible differentiable cells whose macroscopic dynamics can be described by* (1) *and a desired ratio r*_d_ ∈ [0, 1], *design a feedback control law u* = *u*(*t, x*), *where x* = [*x*_1_, ..., *x*_*N*_(*t*)]^┬^, *such that at steady state the consortium is divided into two cell groups whose ratio converges to some desired value, r*_d_, *that is*,

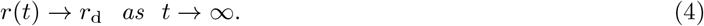

The previous statement can be also reformulated in terms of the control error signal *e*(*t*) ≔ *r*_d_ − *r*(*t*), by requiring that it goes to zero at steady state, that is, *e*_∞_ = 0, where *e*_∞_ ≔ lim_*t*→∞_ *e*(*t*).

Notice that the ratiometric control problem as defined in (4) can only be solved if all cells in the consortium are controllable, as described in Section 2.1.

We present here two different feedback control strategies to solve the ratiometric control problem (see Figure 3), an on-off relay controller and a proportional-integral (PI) controller. Both solutions are easy to implement, robust, and often used in other control applications of cell populations in microfluidic devices [25–28] (see STAR Methods 7.4).

**Figure 3:**
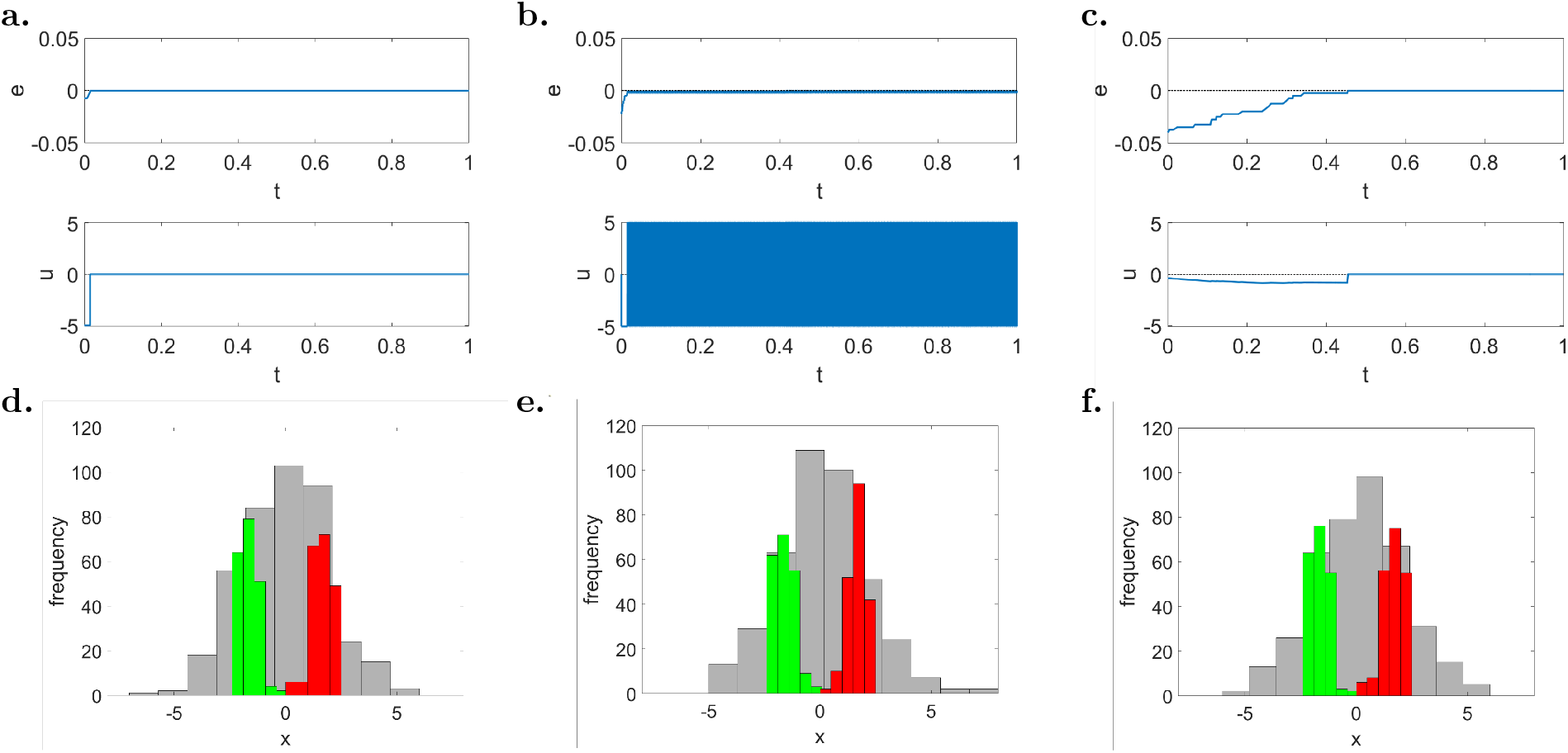
Feedback control strategies are effective to balance two groups of controllable reversible cells to 1:1 ratio (*r*_d_ = 0.5). (**a-c**) Evolution of the error signal *e*(*t*) and of the control input *u*(*t*) for (**a**) the first implementation of the relay controller (22), (**b**) the second implementation of the relay controller (24), (**c**) the PI controller (26)-(27). (**d-f**) Distribution of the cells state at the beginning of the simulation (*t* = 0 a.u., gray histogram) and at steady state (*t* = 1 a.u., green and red histograms), for (**d**) the first implementation of the relay controller, (**e**) the second implementation of the relay controller, (**f**) the PI controller. The green and red bars in panels **(d-f)** correspond to cells being in the basin of attraction of *A*_*i*_ and *B*_*i*_, respectively. The maximum control input is set to *ū* = 5 and the gains of the PI controller are set to *k*_P_ = 30 and *k*_I_ = 10. All cells (*N* = 400) have initial conditions *x*_*i*_(0) drawn from the normal random distribution 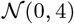, and the parameters *η*_*i*_ are drawn with uniform distribution from the interval [1, 5], therefore all cells are controllable, as no monostable 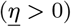 and no unswitchable 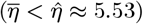 cells are present in the population. (See also Supplemental Figure 8 for more simulations with different desired ratios *r*_d_.)

*Relay controllers* (also known as bang-bang controllers) are simple yet effective feedback control laws that, by comparing the current output of the process of interest with its desired value, generate a piece-wise constant input signal *u*_r_(*t*) whose value belongs to a discrete set 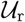. Here, we propose the use of two alternative implementations of the relay controller, the former where the control input can also be set to zero, i.e., 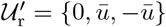, and the latter where *u*_r_ is always non-zero, i.e., 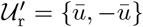.

In the ideal case where all cells are controllable, the first implementation of the relay controller guarantees finite time convergence to zero of the error signal (Figure 3.a). The second implementation instead can only guarantee bounded convergence of the error to zero since the input signal cannot be turned off once the error reaches zero. Hence, when such an implementation is adopted, the control input will continue to oscillate between its possible values (Figure 3.b). As is common practice in applications where noise and uncertainties are unavoidable, a dead-zone or a delay can be added in the control loop to avoid high frequency oscillations of the control input that may cause excessive stress to cells and to the actuation system [29]. The details of the proof of convergence for the proposed relay controllers is reported in STAR Methods 7.5.

An alternative strategy is the use of a *PI controller* that generates a control input *u*_PI_(*t*) computed as the sum of one term proportional (P) to the error *e*(*t*) and an other proportional to its integral (I) in time. In general, PI controllers guarantee zero regulation error at steady state in the presence of constant output disturbances [30]. In our implementation, this controller is complemented with a (anti-windup) reset condition that sets to zero the internal state of the integrator whenever the error signal *e*(*t*) is equal to 0 or changes its sign (see STAR Methods 7.4). When all cells are controllable, this strategy was also proved to solve the ratiometric control problem and guarantee convergence of the error to zero (STAR Methods 7.5).

The evolution of the error signal *e*(*t*) under the action of the PI controller is reported in Figure 3.c. The error converges to zero as expected and the control input *u*_PI_(*t*) also settles to zero after a short transient, similarly to what observed in the first implementation of the relay controller presented before.

Effective balancing of groups of reversible cells is also achieved by feedback control when the goal is to achieve groups of different sizes, that is, for *r*_d_ different from 0.5, e.g., equal to 0.75 or 0.25, corresponding to 1:3 and 3:1 ratios, respectively (Supplemental Figure 8).

### 2.3 Robust bounded regulation of the ratio is still possible in the presence of cell variability and physical constraints

When uncontrollable cells are present in the consortium, that is, cells that cannot be moved from one group to the other in response to any admissible inputs, the ratiometric control problem cannot be solved asymptotically, that is, we cannot guarantee that *e*_∞_ → 0 for an arbitrary initial configuration of the consortium. However, we can still guarantee that the absolute value of the steady-state error |*e*_∞_| will be upper bounded by some positive quantity *e*_*r*_, that is, |*e*_∞_| ≤ *e*_*r*_. This effect is well illustrated in Figure 4, where it is shown that, regardless of the control algorithm being used, the error *e*(*t*) approaches, but does not converge exactly to, zero. The error bound at steady state will depend upon the interplay between heterogeneity of the cells dynamics, and the constraints on the maximum input value of *ū* that can be applied to the cells, as discussed in Section 2.1,.

**Figure 4:**
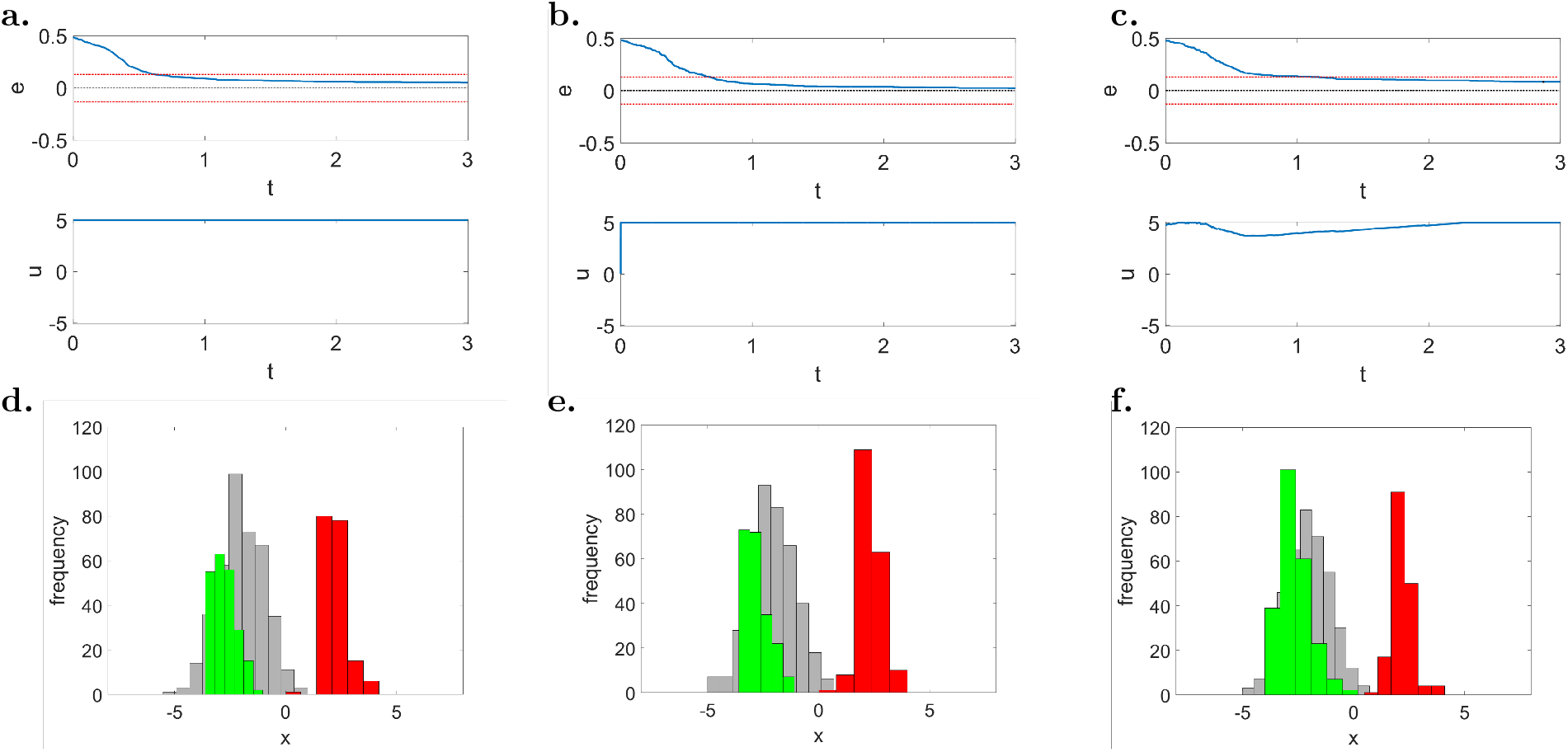
Balance to 1:1 ratio is achieved with a residual steady-state error in the presence of uncontrollable cells (*r*_d_ = 0.5). (**a-c**) Evolution of the error signal *e*(*t*) and of the control input *u*(*t*) for (**a**) the first implementation of the relay controller (22), (**b**) the second implementation of the relay controller (24), (**c**) the PI controller (26)-(27). (**d-f**) Distribution of the cells’ state at the beginning of the simulation (*t* = 0 a.u., gray histogram) and at steady state (*t* = 3.0 a.u., green and red histograms), for (**d**) the first implementation of the relay controller, (**e**) the second implementation of the relay controller, (**f**) the PI controller. The green and red bars in panels **d-f** correspond to cells being in the basin of attraction of *A*_*i*_ and *B*_*i*_, respectively. The maximum control input is set to *ū* = 5 and the gains of the PI controller are set to *k*_P_ = 30 and *k*_I_ = 10. All cells (*N* = 400) have initial conditions *x*_*i*_(0) drawn from the normal random distribution 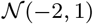, and the parameters *η*_*i*_ are drawn with uniform distribution from the interval [−1, 14], therefore both monostable 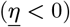 and unswitchable 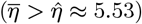 cells can be present in the population. The steady-state errors observed in the *in silico* experiment are equal to **(a)** 0.05, **(b)** 0.0225, and **(c)** 0.0825. Note that all the observed errors are below the theoretical upper bound on the control error that can be estimated using (5) as 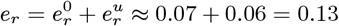 (depicted in the panels **a-c** as red dashed lines). (See also Supplemental Figure 9 for more simulations with different desired ratios *r*_d_.)

For higher number of cells, the upper bound *e*_*r*_ can be estimated as being composed by two terms, that is,

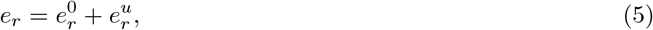

each related to the probability of finding one of the two types of uncontrollable cells (i.e., monostable and unswitchable, respectively) in the consortium (Figure 2). The first term, denoted as 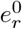, is related to the fraction of monostable cells, associated to a non-positive value of *η*_*i*_, and so admitting only one stable equilibrium point for all values of *u*. Assuming that the probability distribution from where the parameters *η*_*i*_ are drawn is known, we can estimate 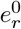 as

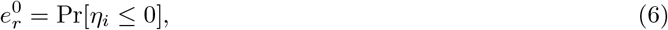

where Pr denotes the probability measure. The second term, denoted as 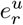, is related to the fraction of unswitchable cells, that is, cells that are bistable but cannot be switched by any admissible value of the control input 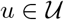. For a given maximum input value *ū*, this fraction of uncontrollable cells can be estimated as the probability that the parameter *η*_*i*_ is greater than 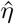, that is, 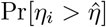. Therefore, the residual error at steady state due to this second type of uncontrollable cells can be quantified as

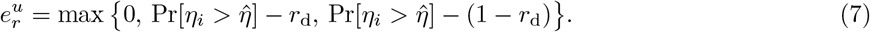

This second residual error term is evaluated by considering the worst case scenario in which unswitchable cells are initially all in one group. Moreover, its value depends on the relationship between the desired ratio for that group (either *r*_d_ or 1 − *r*_d_) and the number of unswitchable cells therein, because they affect the error only when this number exceeds the desired value (see STAR Methods 7.6 for details).

Similar results are observed also when the desired goal is to split the cell population into groups of different sizes, e.g., with 3:1 or 1:3 ratios (Supplemental Figure 9), confirming that this undesired effect is not due to the particular control strategy adopted or to the chosen desired ratio.

### 2.4 Ratiometric control enables cooperative bioproduction in microbial consortia

So far the analysis has been conducted by considering the scalar model (1) capturing the macroscopic bistable nature of the cells considered in this paper. As we are going to show by means of the representative application that follows, the behavior captured by the reduced model in (1) is qualitatively preserved also in more complex and realistic cell models exhibiting the required memory-like property. Therefore, albeit simple, we demonstrate that the model in (1) can be effectively used to design feedback control laws to solve the ratiometric control problem in realistic applications.

As a representative case of study, we consider the agent-based *in silico* implementation of ratiometric control for the bioproduction of protein dimers in microfluidic devices (Figure 5). In this scenario, according to its state (*A* or *B*), each cell in the consortium produces either one of two monomers. By acting on the available control inputs, we want to regulate the relative number of cells producing the two monomers so as to balance the overall production of the resulting dimer.

**Figure 5:**
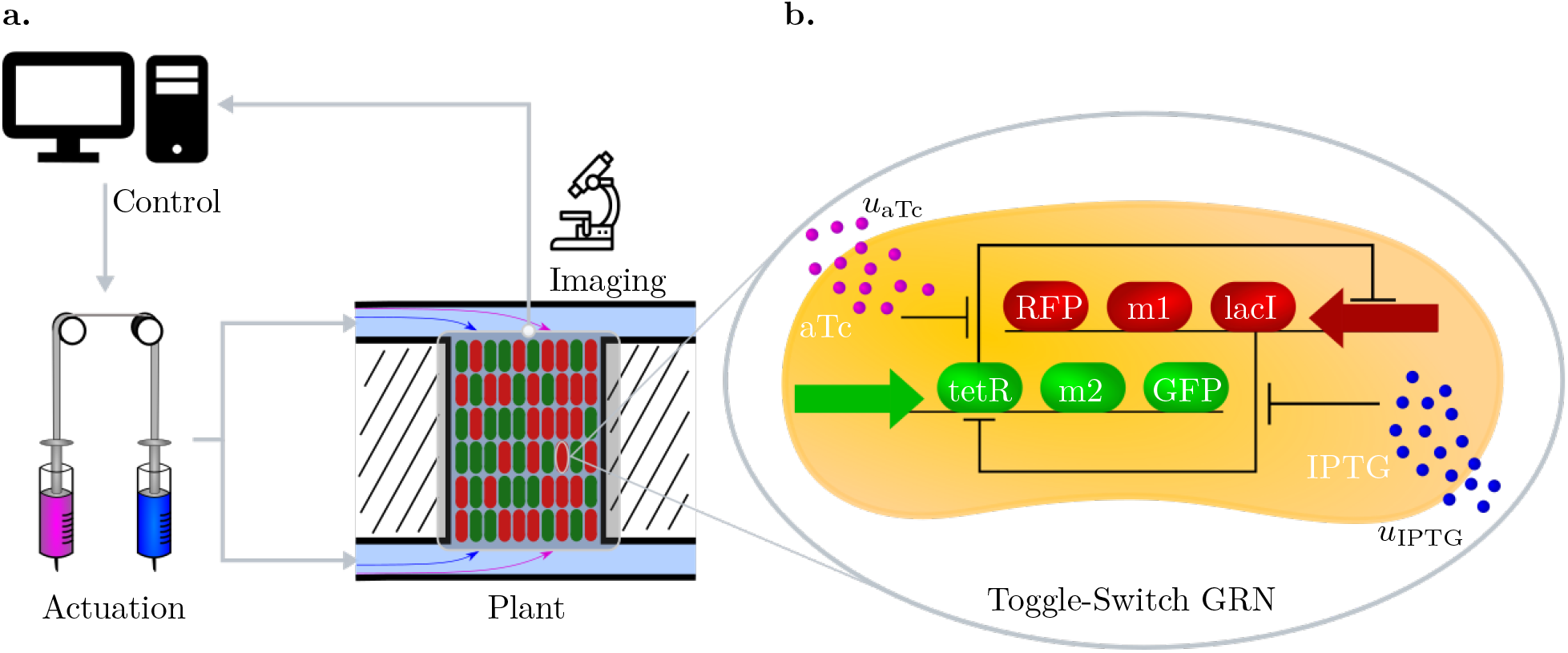
Controlled cooperative bioproduction of a dimer in microfluidic devices. (**a**) Reversible differentiable cells are hosted in microfluidic chambers where they grow and produce a specific molecule related to the corresponding active state of their internal memory. The current ratio of the two cell groups in the chamber, and hence the production level of the corresponding monomers, is evaluated by measuring with a fluorescence microscope the expression of reporter proteins in each cell. This information is acquired by the feedback control algorithm that compares the current ratio *r*(*t*) to the desired ratio *r*_d_ and computes online the corresponding control inputs. Finally, these signals are sent to the cells by actuating a pair of syringes connected to the microfluidic chambers and containing mixtures of growth medium and inducer molecules. (**b**) The required reversible bistable memory mechanism is implemented by using an inducible toggle-switch. Depending on which of the two repressor proteins, either LacI or TetR, is currently expressed, the cell produces the corresponding monomer and the reporter protein (either M1 and RFP or M2 and GFP, respectively). The state of the toggle-switch can be flipped by changing the concentration of the inducer molecules aTc and IPTG in the microfluidic chamber (denoted as *u*_aTc_ and *u*_IPTG_), which diffuse through the cell membrane and bind to TetR and LacI, respectively.

We assume that the mechanism required by the *E.coli* cells to guarantee their correct coordinated behavior is implemented by means of an inducible genetic toggle-switch [31]. Specifically, we consider the circuits design presented in [27] and further analyzed in [32–35]. This genetic regulatory network consists of two repressor proteins, LacI and TetR, both repressing each other’s promoter, so that only one protein is fully expressed at any time. The expression level of the two repressor proteins can be flipped by changing the concentration of two inducer molecules, aTc and IPTG. The former input, aTc, binds to TetR, increasing the rate of production of LacI, and therefore causing the cell to converge to the steady-state corresponding to high expression of LacI. Analogously, IPTG binds to LacI, causing the commitment of the cell to a steady-state corresponding to high expression of TetR.

The dynamical model of each cell is described in details in STAR Methods 7.7, in which the variables *u*_aTc_ and *u*_IPTG_ (as reported in Figure 5) denote the concentrations of the inducer molecules in the growth medium of the microfluidic chambers and they represent the control inputs that can be applied to all cells to change their production role in the consortium.

We further assume that the genes m1 and m2 encoding the two monomers of interest are each fused to one of the repressor genes lacI and tetR of the toggle-switch circuit. So that, at steady state, each cell fully produces only one monomer at the time and at a rate assumed to be proportional to the concentration of the corresponding repressor protein. Reporter genes of red and green fluorescent proteins (RFP and GFP) are also bound to the repressor genes to monitor the current level of production of the monomers by using fluorescence microscopy (Figure 5). Finally, we assume that the two monomers have equal transcription and translation rates. Therefore, for the dimer to be produced at high rate, the consortium must be split and maintained into two symmetric groups with a 1:1 ratio, that is, we set *r*_d_ = 0.5.

In the *in silico* experiments, we also take into account realistic physical and technological constraints of a possible implementation in the microfluidic experimental platform described in [11, 25]. Specifically, we consider constraints on (i) the possible classes of input signals that can be generated by the actuators, (ii) an upper bound on the switching frequency of the inputs to limit osmotic stress to the cells, (iii) a time delay accounting for the time the chemical inducers take to flow from the reservoirs to the cell chambers, and (iv) a safety lower bound on the sampling time of the measurements to avoid excessive photo-toxicity (see STAR Methods 7.8).

By using a timescale separation argument (see STAR Methods 7.7), it is possible to capture the cell dynamics using the following reduced-order model

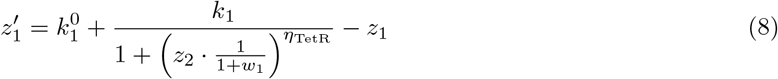

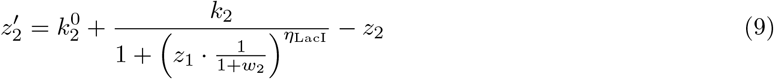

where the symbol (′) denotes the derivative with respect to rescaled time *t′*,

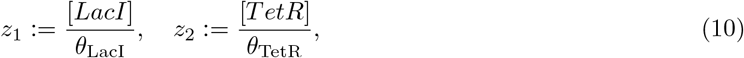

and

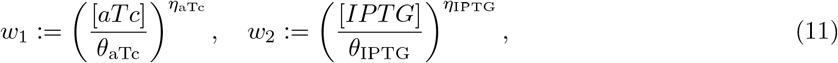

where [·] denotes concentrations.

It is possible to verify that system (8)-(9) is memory-like as required, since in its domain of definition the phase portrait is qualitatively similar to that of the system in (14a) (Figure 7.a). Moreover, this dynamical system has also been numerically investigated in [27, Supplementary Figure 6] where it has been shown that its dynamics exhibit bistable behavior with hysteresis.

*In silico* control experiments have been conducted by using *ad hoc* implementations in BSim [22, 23] of the two feedback control algorithms presented in Section 2.2 (see STAR Methods 7.9). To test the relay control strategy, we assumed that the actuation of the inputs is realized using an ordinary T-junction [36], which allows only one inducer molecule at a time to be injected into the microfluidic chambers. For the PI controller we assumed that the actuation is realized by a Dial-A-Wave system, as described in [37]. This actuation system is more advanced than the previous one as it allows mixtures of the two inducers to be injected in different proportions into the chambers. Both feedback control algorithms take into account the characteristics of the experimental platform and in particular of the actuators. Full details about the control algorithms and the technological constraints of the platform are reported in STAR Methods 7.9.

The agent-based simulations in BSim accurately capture cells’ reproduction, spatial distribution and geometry of the cells and of the microfluidic chambers, diffusion of chemicals into the environment and, more importantly, flush-out of the cells from the chambers. Further details on the stochastic simulation algorithm, geometry and other parameters used for *in silico* experiments in BSim are reported in STAR Methods 7.8.

We observed that both controllers can successfully regulate the populations’ ratio to the desired value after relatively short transients (Figure 6 and Supplemental Video 1). The relay controller shows a faster response with more severe oscillations (Figure 6.a-c), while the PI controller presents a smoother but slower response with higher accuracy at steady state (Figure 6.d-f). This is expected as it is well-known that the relay control strategy is in general more robust to uncertainties and noise affecting the controlled process but has poorer accuracy at steady state; the PI control strategy showing better steady-state performance thanks to the presence of an integral action.

**Figure 6:**
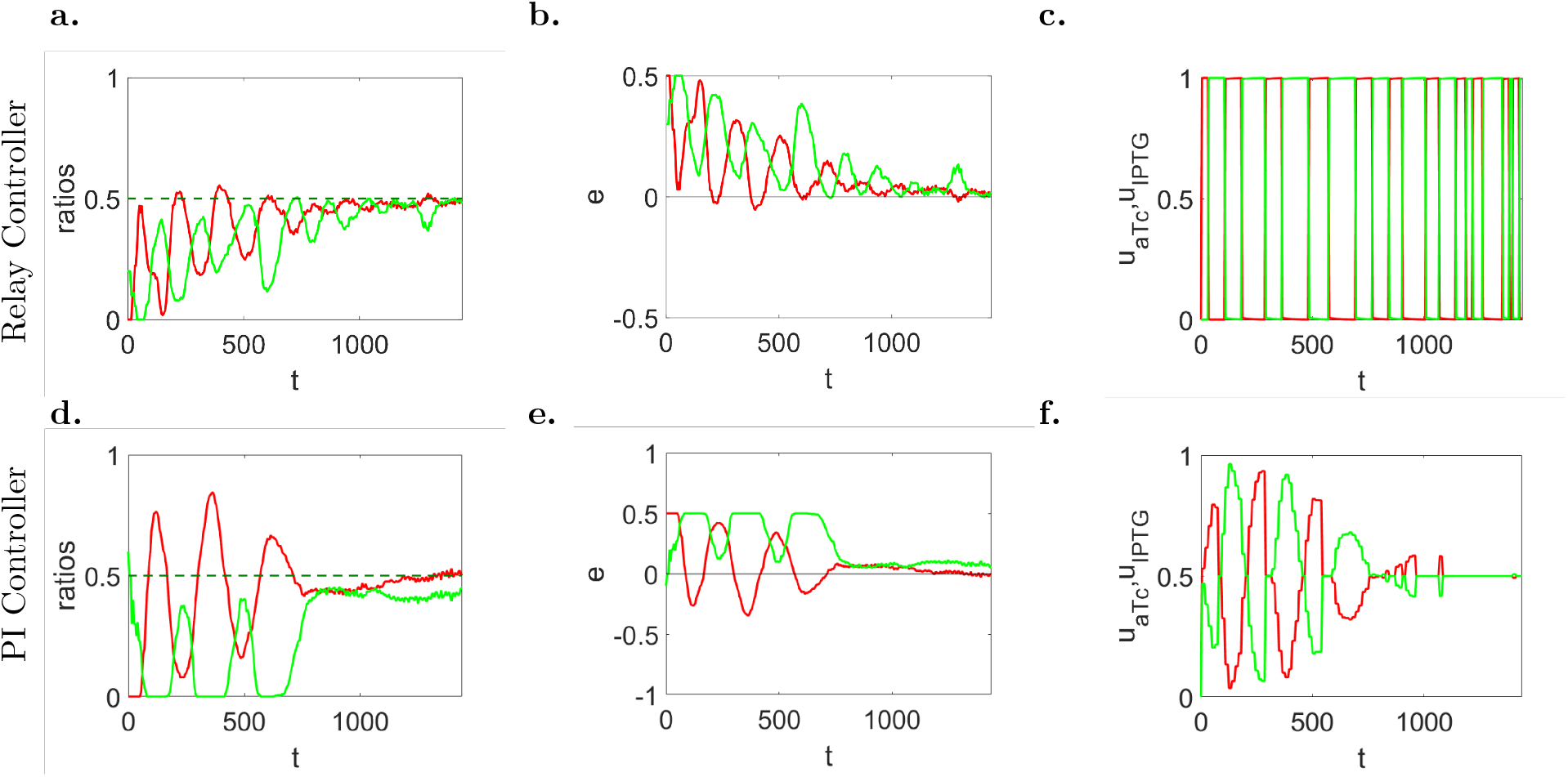
Cooperative production of two monomers to 1:1 population ratio can be achieved by means of feedback ratiometric controllers in microfluidics. (**a,d**) Evolution in time of populations’ ratio *r*_*A*_ (solid green line) and *r*_*B*_ (solid green line) with their respective desired reference values in dashed lines (*r*_d_ = 0.5), (**b,e**) of the error signals *e*_*A*_ (solid green line) and *e*_*B*_ (solid red line), and (**c,f**) inducer control signals *U*_aTc_ (solid red line) and *U*_IPTG_ (solid green line), normalized to their maximum values *u*_aTc_ and *u*_IPTG_, respectively. (**a-c**) Parameters of the relay control (55): *U*_aTc_ = 60 ng/mL, *U*_IPTG_ = 0.5 mM. (**d-f**) Parameters of the PI controller (56): *U*_aTc_ = 100 ng/mL, *U*_IPTG_ = 1 mM, *k*_P,1_ = 100, *k*_P,2_ = 1.5, *k*_I,1_ = 1.5, *k*_I,2_ = 0.05. Cells (about 200) in the simulated microfluidic chamber (with dimensions 40 *μ*m × 50 *μ*m × 1 *μ*m) have all the same parameters’ value, and their evolution has been obtained using the agent-based simulator BSim [22, 23] (See STAR Methods 7.8 for further details on the simulator setup). (See also Supplemental Video 1, and Supplemental Figure 10 for more *in silico* experiments with different desired ratios *r*_d_.)

The difference in the performance of the two strategies is also stressed by the different actuation systems employed in our experiments. Indeed, the Dial-A-Wave system allows for a finer regulation of the concentrations of the inducer molecules than the simpler (and cheaper) T-junction, allowing better accuracy of the control system at steady state. Similar performances are obtained also when the goal is changed to achieve different population ratios, e.g., a 1:3 ratio or a 3:1 ratio (Supplemental Figure 10). These scenarios may correspond, for example, to situations in which the two monomers have equal transcription rates but different translation rates, requiring the consortium to be split into two asymmetric groups, for efficient production of the dimer.

Besides biochemical noise, the fluctuations at steady state (Figure 6.a,d) are essentially due to cells being flushed out of the microfluidic chamber as they grow and duplicate. This effect is more relevant when cells are host in a small chamber and becomes less significant as the size of the growth chambers increases. Indeed, for the sake of simplicity, assuming the chamber to be a square of side *ℓ*, the magnitude *ε* of the fluctuations is proportional to 1/*ℓ* (see STAR Methods 7.10), hence the fluctuations increase as the chamber size decreases (see Supplemental Figure 11 and Supplemental Video 2), and vice versa. Finally, the ratiometric controllers presented here also showed robustness to cell-to-cell variability in their response, as shown in Supplemental Figure 12.

## 3 Discussion

We presented a general framework to guide the design of external feedback controllers for labor division in microbial consortia. We showed that, by exploiting the memory-like property of reversible differentiable cells, a single-strain cell population can be divided into two groups, expressing different sets of genes, whose relative numbers, i.e., the ratio, can be regulated by means of common exogenous inputs. We showed by means of a representative example that ratiometric feedback controllers can robustly stabilize a cell population, endowed with a genetic toggle-switch functioning as a bistable memory, and can guarantee the balance between the two groups of cells even in the presence of realistic physical and technological constraints of the experimental microfluidic platform we considered.

A fundamental open problem in multicellular control applications is to guarantee coexistence of different microbial strains growing in the same environment. Although some solutions were proposed in the literature that rely on the use of additional genetic pathways embedded into the cells, the ratiometric control framework we presented here provides an alternative approach that might be more appropriate in other scenarios, for example, in industrial applications, where efficient production is strongly required. However, the scaling up of the ratiometric control strategies we proposed here for the balancing of microbial communities in bioreactors is still not fully investigated; in previous work [19] we only considered the problem of regulating the relative number of two *different* microbial strains, while the case of a monostrain consortium composed by reversible cells has not yet been explored and is left for future work.

Finally, we wish to highlight that ratiometric control of a population of reversible cells by means of a common input signal is only made possible by the heterogeneity of their response to that input. Indeed, heterogenous reversible cells characterized by different parameter values switch at different time instants when subject to the same input, and this is a crucial property that allows their state to be controlled by an external feedback action. Stochastic effects, such as biochemical noise or delays, by amplifying the cell-to-cell variability can indeed facilitate the stabilization of the reversible cells into different groups, as it was demonstrated in the *in silico* experiments we provided. A more in-depth analytical investigation of their effect is left for future work.

## 4 Acknowledgments

The authors wish to thank Prof. S. John Hogan, University of Bristol, U.K. for his insightful comments on the convergence of discontinuous maps, and also Dr. Lorena Postiglione, University of Bristol, U.K. for her advice about experimental microfluidic platforms.

M.d.B. and D.F. wish to acknowledge support from the European Union’s Horizon 2020 research and innovation programme under grant agreement No 766840 (COSY-BIO). D.F. also wishes to acknowledge support from the research grant “BIOMASS” funded by the University of Naples Federico II - “Finanziamento della Ricerca di Ateneo (FRA) - Linea B”.

## 5 Author Contributions

D.F. and M.d.B. designed the research; D.F. and D.S. designed the controllers and carried out the modeling; D.S. implemented the controllers in BSim and carried out the *in silico* experiments; D.S. and D.F. developed the analysis; M.d.B. supervised the project; D.S., D.F. and M.d.B. analyzed the data and wrote the manuscript.

## 6 Declaration of Interests

The authors declare no competing interests.

## 7 STAR Methods

### Lead contact and materials

Further information and requests for resources should be directed to and will be fulfilled by the Lead Contact, Mario di Bernardo.

### Methods Details

#### 7.1 Modeling reversible differentiable cells

We assume that the macroscopic behavior of any cell in the consortium we wish to control can be modeled in the domain of interest by a dynamical system of the form

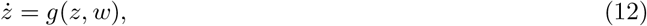

with 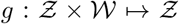 being a smooth vector field, 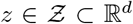 the state variables and 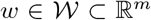 the exogenous input variables, representing, for instance, the concentrations of chemical species inside the cell and those of control inducer molecules into the environment, respectively. We assume that stochastic effects, such as fluctuations due to biochemical reactions, do not significantly alter the behavior of the system at steady state; that is, the region of attraction of any stable equilibrium point of the dynamical system 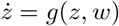 is large enough so that stochastic noise does not cause undesired switch from one equilibrium point to another.

As discussed in the Introduction and in Section 2.1, we are interested to robustly regulate the behavior of reversible cells, in particular we focus our attention to a specific class of cells whose dynamics satisfy the following fundamental property (Supplemental Figure 7.a).

##### Definition

*Consider a dynamical system of the form*

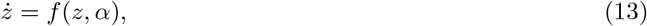

*where* 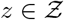 *and the parameter* 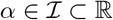 *depends on some exogenous input signal* 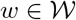, *that is, α = α(w). We say that system* (13) *is* memory-like *if for all* 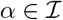 *it is globally topologically equivalent, in its domain of definition* 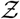, *to the dynamical system*

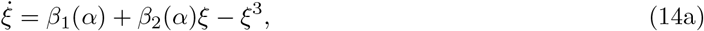

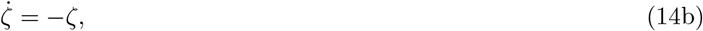

*with* 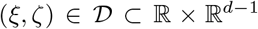, *β*_1_ *and β*_2_ *being in bijective correspondence with α, that is, there exists a homeomorphism* 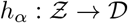 *mapping orbits of the first system onto orbits of the second system, preserving the direction of time. Moreover, β*_1_ *and β*_2_ *are monotonic functions with respect to α, and β*_1_(0) = 0.

As state variables *ξ* and *ζ* in system (14a)-(14b) are uncoupled and *ζ*(*t*) asymptotically converges to 0 for all initial condition *ζ*(0), the qualitative behavior of the system can be completely described by the dynamics of *ξ*. Note that, (14a) is the topological normal form of the so-called cusp bifurcation [38, p.301], which can exhibit hysteretic behavior.

Under the assumption that system (12) is memory-like, hence being topological equivalent to system (14a), and by further assuming that (i) the value of *β*_*i*,1_ is the same across the entire population, i.e., *β*_*i*,1_ = *u*, ∀*i*, and that (ii) *β*_*i*,2_ is constant but different for each cell *i*, i.e., *β*_*i*,2_ = *η*_*i*_, any reversible cell in the consortium can be modeled as in (1), where *u* is an *equivalent input* signal that depends on *w*. However, it can be showed that the previous assumptions, albeit simple, lead to more conservative results for the original model (13).

Indeed, by defining 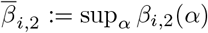 and 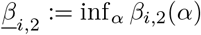, a cell *i* whose dynamics is described by model (13), is controllable (see Section 2.1) if (i) 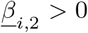, and (ii) 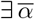 such that 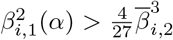, for all 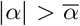. The first condition implies that the cell is not monostable, that is, the system has two stable equilibrium points for *α* = 0. On the other hand, having assumed that *β*_*i*,1_ and *β*_*i*,2_ are monotonic functions with respect to *α*, the second condition implies that the cell is controllable, that is, there exists a value of the input *α* such that the bistable system becomes monostable (see STAR Methods 7.2).

Finally, by choosing the parameter *η*_*i*_ in model (1) as

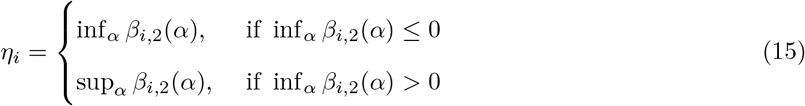

and by setting *ū* = sup_*i,α*_|*β*_*i*,1_(*α*)|, the cells described by model (1) that are controllable are indeed a smaller subset of those described by model (13), that is, by making this choice for the parameters we are considering a worst-case scenario. Thus, all the results presented for model (1), included the proof of convergence of the proposed controllers, also hold for model (13).

#### 7.2 Derivation of 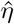

In this section we show how the expression of 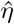 in (2) was derived. The quantity 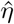 represents the limit value of *η*_*i*_ such that the equation 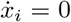 with *η*_*i*_ > 0 and *u* = *ū*, where 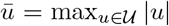, has a unique solution. That is, any cell whose behavior can be described by (1) with 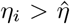, cannot be switched from *A*_*i*_ to *B*_*i*_, or vice versa, by any input value 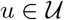, because it always admits at least two solutions and it is therefore uncontrollable.

To this aim, we have to characterize the bifurcation points at which two equilibrium points of (1) coalesce and disappear [38], under variations of the control input *u* and of the parameters *η*_*i*_. The position of the limit points can be easily obtained as:

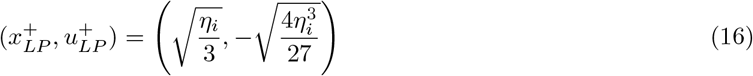

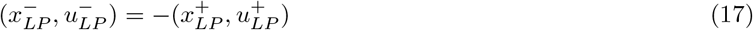

Hence, equation 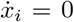 admits at least two solutions if its parameters and the control input satisfy the relationship

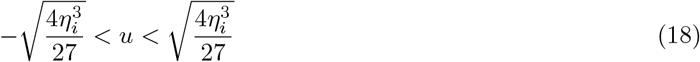

Therefore, it is easy to verify from (18) that, given the maximum value of the control input *ū*, the threshold value 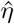 over which a cell becomes uncontrollable is

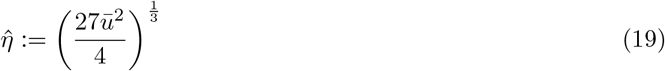

#### 7.3 Event-driven modeling of the control error evolution

Here we derive an event-driven model for the evolution of the control error *e*(*t*) = *r*_d_ − *r*(*t*), presented in Section 2.2. Recall that the finite set of all cells in the consortium and its cardinality at time *t* are denoted by 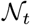 and *N*(*t*), respectively. Now, we denote with 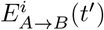 the event at time instant *t* = *t′*, corresponding to when the state *x*_*i*_ of cell 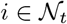 enters the basin of attraction of the equilibrium point *B*_*i*_, that is, 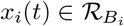 for *t* ∈ [*t′, t′* + *ε*], and 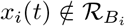 for *t* ∈ [*t′* − *ε, t′*), where *ε* is a small positive real number. Likewise, we denote by 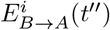 the event at time instant *t* = *t′′*, corresponding to when the state of cell 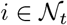 enters the basin of attraction of *A*_*i*_. Specifically, for solutions to dynamical system (1) with *u* = 0 and *η*_*i*_ > 0, an event 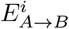 occurs when the state of cell *i*, *x*_*i*_, crosses zero and becomes positive, while an event 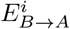 occurs when *x*_*i*_ becomes negative. In this case the threshold at zero is defined by the unstable equilibrium point at the origin dividing the regions of attraction of *A*_*i*_ and *B*_*i*_. Moreover, we denote by 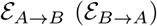 the sets of all events 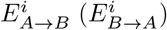 occurring for all *i* at any time *t*, and we denote by 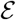 the set of all events occurring in the population, that is, 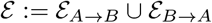.

To derive the discrete (event-driven) dynamics of the control error *e*(*t*), we make the following standing assumptions:

##### Assumption 1

*At any time t only one event in* 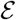 *can occur, that is, there exists a unique* 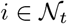 *such that either* 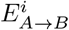 *or* 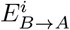, *but not both, occurs at time t.*

##### Assumption 2

*The number of cells in the host chamber is assumed to be constant, that is, N*(*t*) = *N, for all t.*

Assumption 1 implies that any two events in 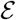 cannot occur simultaneously. Assumption 2 follows from the fact that, after a short transient from the beginning of the experiment, cells grow occupying the entire host chamber. From this time on, cells exceeding the maximum capacity of the chamber are flushed out. Therefore, the number of cells in the chamber can be assumed with a good approximation to be constant (except for a small, negligible oscillation due to flush-out, further discussed in STAR Methods 7.10).

Then, there exists a sequence of discrete time instants {*t*_*k*_}_*k*∈ℕ_, each one corresponding to the occurrence of an event in 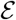 and such that *t*_*k*+1_ = *t*_*k*_ + Δ*t*_*k*_, where Δ*t*_*k*_ > 0 is the time interval between two consecutive events occurring at time *t*_*k*_ and *t*_*k*+1_. respectively. Moreover, from Assumption 2 it follows that the functions *n*_*A*_(*t*) and *n*_*B*_(*t*), defining the number of cells converging to either *A*_*i*_ or *B*_*i*_, respectively, are piece-wise constant functions, that is, *n*_*A*_(*t*) = *n*_*A*_(*t*_*k*_), and *n*_*B*_(*t*) = *n*_*B*_(*t*_*k*_), ∀*t* ∈ [*t*_*k*_, *t*_*k*+1_). Since *n*_*A*_(*t*) and *n*_*B*_(*t*) are constrained by the relation *n*_*A*_(*t*) + *n*_*B*_(*t*) = *N*, for all *t*, for the sake of brevity, we will refer to *n*(*t*) ≔ *n*_*B*_(*t*) only; *n*_*A*_(*t*) being given by *N* − *n*_*B*_(*t*).

That said, being *n*(*t*) the number of cells in the basin of attraction of *B*_*i*_ at time *t*, we can write the following discrete-time update law:

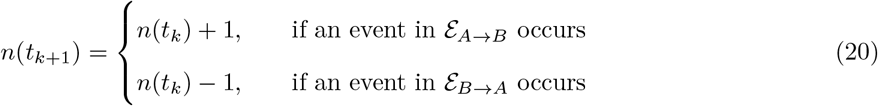

As a consequence, since 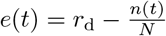, we have that when an event in 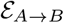 occurs, then

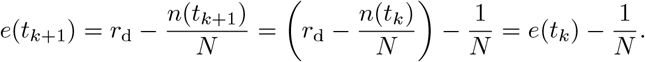

A similar reasoning holds when an event in 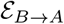 occurs. Therefore, the discrete-time dynamics of the control error can be expressed as:

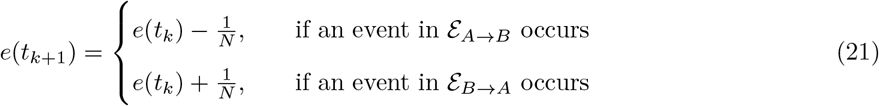

#### 7.4 Implementation details of the proposed feedback control algorithms

Here we discuss the design of the control strategies proposed to solve the ratiometric control problem, namely, the relay controller and the PI controller.

##### Relay control algorithm

A *relay controller* is a feedback control law that generates a piece-wise constant input *u*_r_(*t*) by comparing an output measured from the plant to some desired reference value *r*_d_. The input *u*_r_(*t*) takes value from a finite set of real values 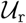, generally composed by only two values, one chosen such that the control error *e*(*t*) = *r*_d_ − *r*(*t*) decreases when *e*(*t*) > 0, and the other such that it increases when *e*(*t*) < 0.

We have considered two implementations of the relay controller. Specifically, the first implementation includes a control shutdown condition when *e*(*t*) = 0, and the second one does not.

Formally, the *first* implementation of the relay control input is defined as:

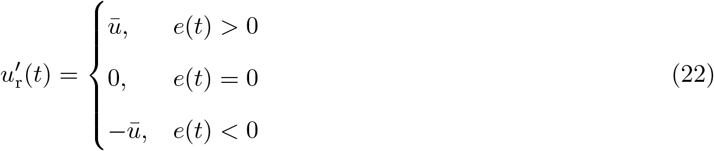

The value *ū* > 0 is chosen such that (ideally) all cells are controllable, that is, for *u′*_r_ = *ū*(*u′*_r_ = −*ū*), the equation 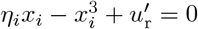 has a unique stable solution, namely 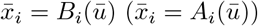, for all *i* such that *η*_*i*_ > 0. This guarantees that when *e*(*t*) > 0 (i.e., *r*_*B*_(*t*) < *r*_d_), the next event occurring must belong to 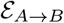, forcing the error to decrease, that is, 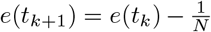. Likewise, when *e*(*t*) < 0 (i.e., *r*_*B*_(*t*) > *r*_d_), *u′*_r_ = −*ū* implies that the next event occurring belongs to 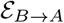, and so 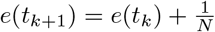. Moreover, notice that, when no control is applied (i.e., *u′*_r_ = 0), each cell will converge to either *A*_*i*_ or *B*_*i*_, depending on its current state, without any other event in 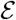 having to occur. Therefore, the shutdown condition ensures that, if there exists a *t*\* such that *e*(*t*\*) = 0, then *e*(*t*) = 0 for all *t* ≥ *t*\*. Combining (22) and (21), the discrete-time update law for the control error becomes:

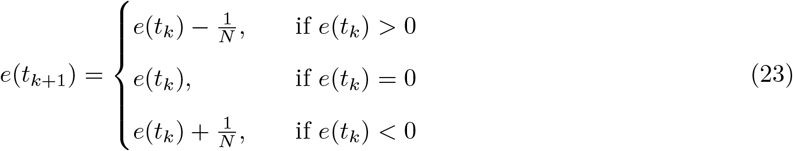

The previous discrete map can also be rewritten as 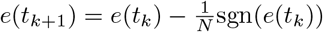.

The *second* implementation of the relay control input, without the shutdown condition, is defined as:

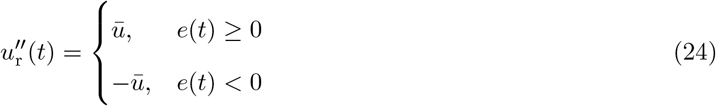

In this case it is not possible to ensure that the error remains equal to zero indefinitely. By using a similar reasoning as before, (21) can be recast as:

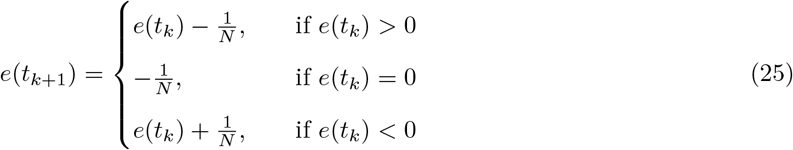

##### PI control algorithm

The control input of the PI controller is defined as:

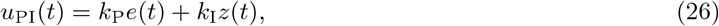

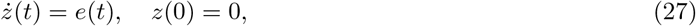

with *k*_P_ and *k*_I_ being positive constants. This control action is complemented with a (anti-windup) reset condition that sets the internal state *z* of the integrator to zero whenever the error becomes 0 or changes its sign. Furthermore, to take into account constraints on the actuation system of the experimental platform, the control input signal *u*_PI_ is assumed to be saturated at *ū* and −*ū*.

The control algorithm guarantees that when *e*(*t*) > 0 the control input *u*_PI_(*t*) is positive and 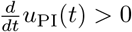. So, *u*_PI_(*t*) will increase and reach some positive value 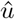 such that, for at least one cell, the equation 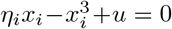 with 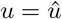 and *η*_*i*_ > 0 has a unique solution, namely 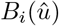 (see Figure 7). The cell will be attracted by this stable equilibrium point and, therefore, there will exist a time instant *t′* such that, for all *t* ≥ *t′*, 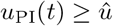 and an event in 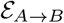 will occur. A similar reasoning holds in the case *e*(*t*) < 0. Hence, it directly follows that the discrete-time update law for the control error *e*(*t*) under the PI control law is the same as in (23).

**Figure 7:**
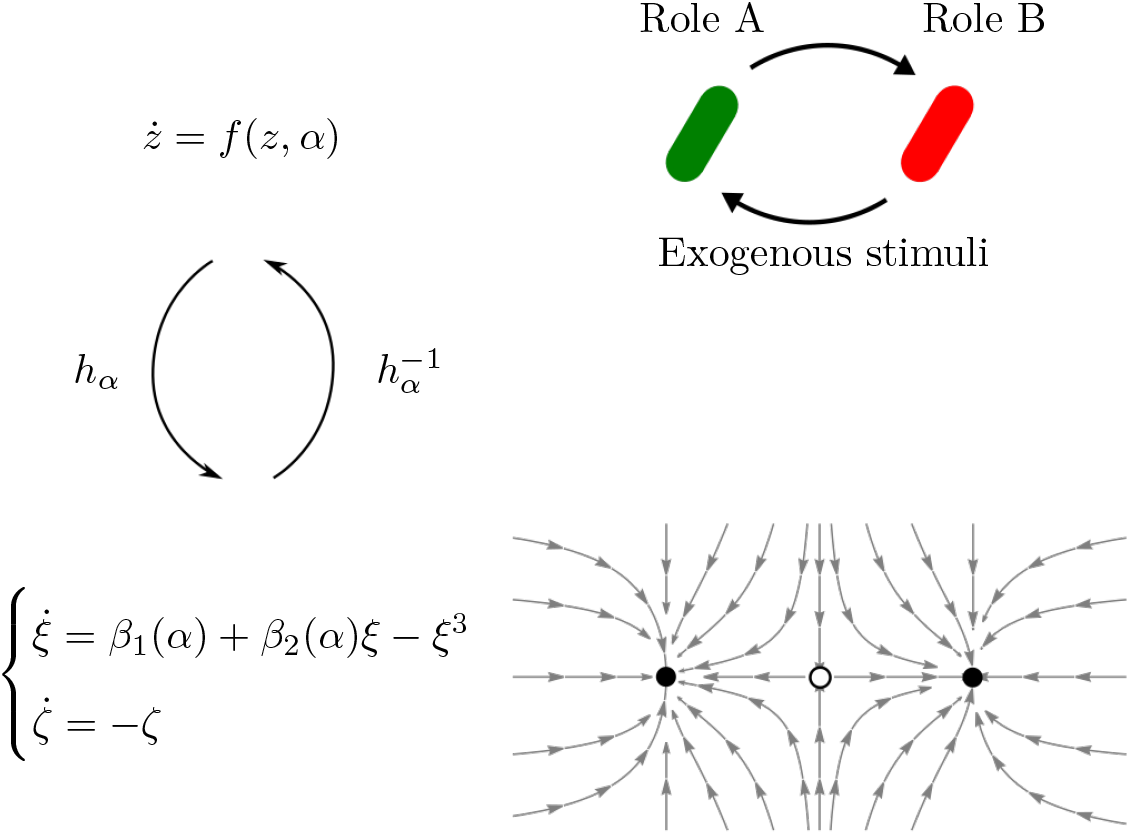
Macroscopic dynamics of reversible differentiable cells with memory-like property can be effectively modeled as a scalar bistable dynamical system. The dynamical system 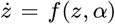 describing the evolution of a reversible differentiable cell with *memory-like* property is topologically equivalent to the system 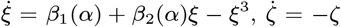. That is, there exists a correspondence between solutions of the two dynamical systems, in particular they have the same number of equilibrium points of the same stability type. Moreover, this equivalence is preserved as the bifurcation parameter *α* varies, to which there correspond some values of the parameters *β*_1_(*α*) and *β*_2_(*α*) in the second system.

#### 7.5 Proof of convergence of the proposed controllers

Here we prove that the feedback control algorithms we proposed ensure convergence to the desired cell ratio and hence solve the ratiometric control problem stated in Section 2.2. To this aim we conduct a convergence analysis on the error map *e*_*k*+1_ = *f* (*e*_*k*_), where *f* : *D* → *D*, with *f* being defined in (23) or (25), and *D* is the domain of definition of the error signal *e*_*k*_. Specifically, we consider some positive definite scalar function *V* : *D* → ℝ, and we prove convergence of *e*_*k*_ to some invariant set by assessing the negative definiteness of the increment function Δ*V* ≔ *V*(*f*(*e*_*k*_)) − *V*(*e*_*k*_) for every point outside this invariant set.

Notice that the domain of definition *D* is *finite*, indeed, from the definition of the control error signal, that is, 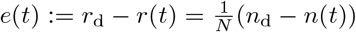, and from the fact that *n*_d_ = *N* · *r*_d_ is an integer constant and *n*(*t*) is integer-valued function taking values in the finite set {0, 1, 2, *..., N*}, we have that

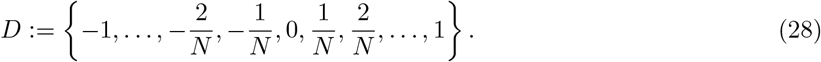

##### Relay control algorithm(22)

To prove convergence of the error signal *e* to 0, let us consider the function *V*(*e*) = *e*^2^. Using the discrete-time map in (23), we have that

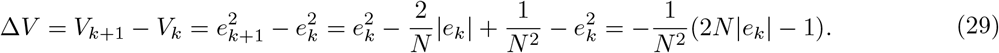

The increment Δ*V* is negative definite everywhere except in the points such that

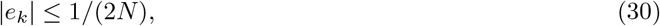

therefore any solution whose initial condition is outside this set converges towards it. Since the only possible value of *e*_*k*_ in *D* satisfying (30) is 0, we can therefore conclude that the error signal *e*_*k*_ converges to 0.

##### Relay control algorithm (24)

From (25) it directly follows that the error dynamics does not admit any fixed point. However, a period-2 cycle exists whose stability we are going to prove by considering the second iterate of the error map in (25):

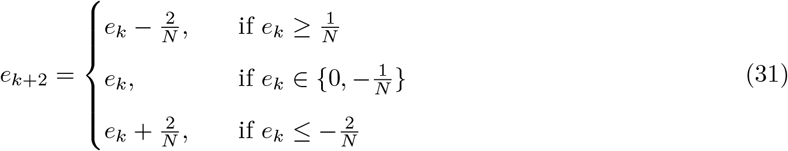

The periodic orbit of the first iterate map (25) corresponds to the pair of fixed points located in 0 and −1/*N* of the second iterate map (31).

Let us consider the function *V*(*e*) = |*e*|. By using the second iterated map in (31) and taking into account (28) we have:

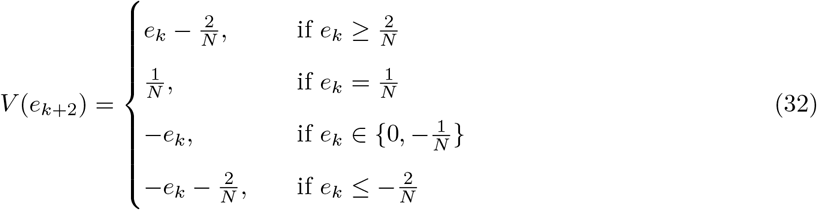

Hence, after some algebra we can write:

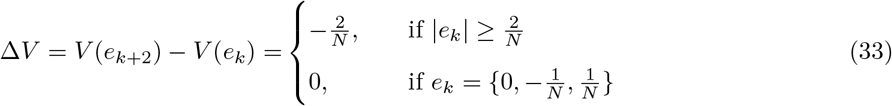

Therefore, we can conclude that all solutions converge to the set where Δ*V* = 0, that is, such that |*e*_*k*_| ≤ 1/*N*, or, equivalently, to the set 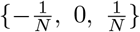. Now, by using the second iterate map (31), we can easily verify that in the set 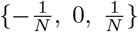 there are two fixed points, namely 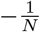 and 0. Therefore, these two fixed points correspond to period-2 cycles for the first iterated map (25). More precisely, both points correspond to the same cycle 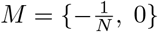, indeed, by applying (25) we have that 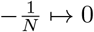 and 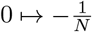. This proves that the error signal globally converges to the period-2 cycle *M*.

##### PI control algorithm (26)-(27)

In this case the discrete-time map describing the dynamics of the control error is also defined by (23). Therefore, the proof of convergence for the PI controller is the same as that of the first implementation of the relay controller (22) presented earlier on.

#### 7.6 Derivation of 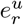

Here we illustrate how the expression of 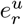 in (7) was obtained. The quantity 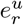 is an upper bound on the residual steady-state error due to the presence of *unswitchable* cells, i.e., bistable cells that cannot be toggled by any admissible value of the control input *u*. The maximum value of this quantity can be estimated by considering the worst case scenarios in which all these uncontrollable cells are initially in one region of attraction and therefore cannot be moved to the other one by any admissible values of *u*.

Without loss of generality, assume that all uncontrollable cells are initially in the region of attraction of their equilibrium point *B*_*i*_ and recall that *r*_d_ denotes the desired ratio of cells converging to *B*_*i*_ at steady state. Denoting with *r*_*u*_ the ratio between the number of uncontrollable cells and the whole cell population, we have that if *r*_*u*_ ≤ *r*_*d*_, that is, the number of uncontrollable cells does not exceed the desired number in *B*_*i*_, then uncontrollable cells do not cause any problem because we do not need to move any of them from where they are. Therefore, in this case, *e*_∞_ = 0. On the other hand, if *r*_*u*_ > *r*_d_, the fraction of uncontrollable cells exceeding the desired ratio, that is, *r*_*u*_ − *r*_d_, will determine an error at steady state, that is, |*e*_∞_| ≤ *r*_*u*_ − r_d_. A similar argument can be stated for the dual scenario in which all uncontrollable cells are initially in the region of attraction of *A*_*i*_. Recalling that in this case we want the ratio of cells converging to *A*_*i*_ to be equal to (1 − *r*_d_), the residual error at steady state in this scenario will satisfy |*e*_∞_| ≤ *r*_*u*_ − (1 − r_d_).

From the previous analysis and exploiting that, for a high number of cells, *r*_*u*_ can be estimated as the probability of *η*_*i*_ begin greater that 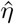, we finally obtain that

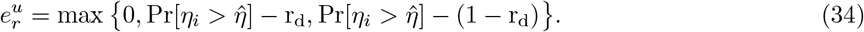

where Pr denotes the probability measure.

#### 7.7 Mathematical model of the inducible toggle-switch

In this section we present the details of the mathematical model used to describe the time evolution of proteins’ concentration in the inducible toggle-switch. The model of the toggle-switch we consider here was originally developed and parameterized from experimental data in [27]. It was further analyzed in [32–35]

The model captures the pseudo-reactions describing transcription

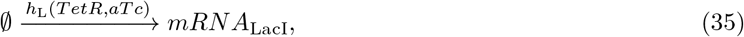

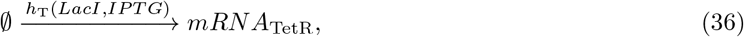

where

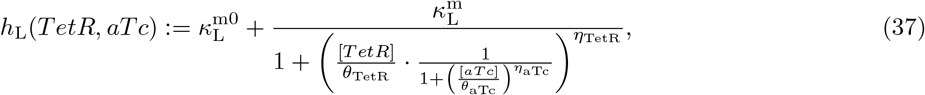

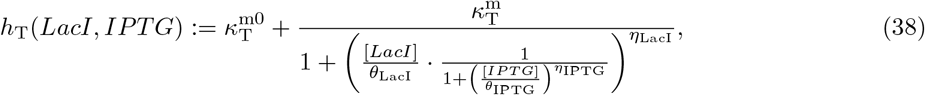

those describing translation

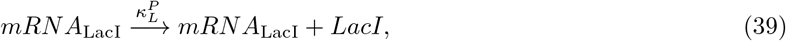

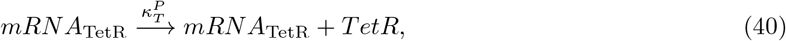

and those related to dilution/degradation

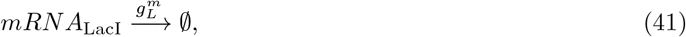

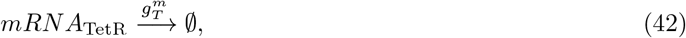

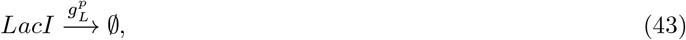

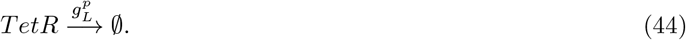

The repressive action of LacI and TetR on each other is captured in (37)-(38) by means of Hill functions (with parameters *θ*s and *η*s). Likewise, the repressions of aTc on TetR and of IPTG on LacI are both described by another Hill function nested in the previous two. In the above pseudo-reactions, the parameters 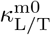, 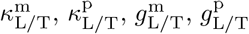 are basal transcription, transcription, translation, mRNA degradation and protein degradation rates, respectively. The value of these parameters are reported in Table 1.

**Table 1:** Values of the parameters of the cell population models (taken from [27]).

|  |  |  |  |
| --- | --- | --- | --- |
| $\kappa_L^{m0}$ | 0.3045 mRNA min<sup>-1</sup> | $g_L^p$ | 0.0165 min<sup>-1</sup> |
| $\kappa_T^{m0}$ | 0.3313 mRNA min<sup>-1</sup> | $g_T^p$ | 0.0165 min<sup>-1</sup> |
| $\kappa_L^m$ | 13.01 mRNA min<sup>-1</sup> | $\theta_{LacI}$ | 124.9 |
| $\kappa_T^m$ | 5.055 mRNA min<sup>-1</sup> | $\eta_{LacI}$ | 2.00 |
| $\kappa_L^p$ | 0.6606 a.u. mRNA<sup>-1</sup> min<sup>-1</sup> | $\theta_{TetR}$ | 76.40 |
| $\kappa_T^p$ | 0.5098 a.u. mRNA<sup>-1</sup> min<sup>-1</sup> | $\eta_{TetR}$ | 2.152 |
| $k_{aTc}$ | 0.04 min<sup>-1</sup> | $\theta_{aTc}$ | 35.98 |
| $k_{IPTG}$ | 0.04 min<sup>-1</sup> | $\eta_{aTc}$ | 2.00 |
| $g_L^m$ | 0.1386 min<sup>-1</sup> | $\theta_{IPTG}$ | 0.2926 |
| $g_T^m$ | 0.1386 min<sup>-1</sup> | $\eta_{IPTG}$ | 2.00 |

The pseudo-reactions listed above can be used to obtain the following *deterministic model* of the inducible toggle-switch:

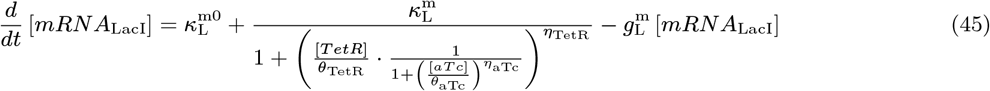

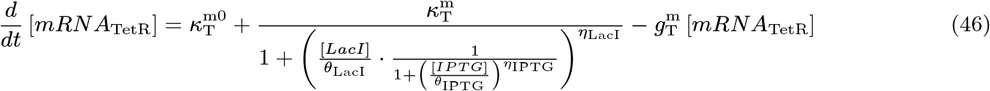

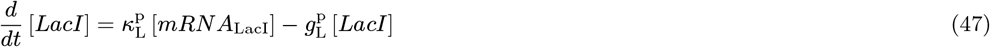

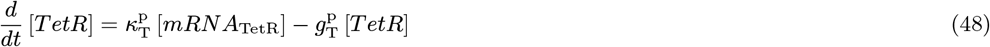

where the quantities in the form [·] denote concentrations of molecules inside each cell.

The model is complemented by also taking into account the diffusion of aTc and IPTG across the cell membrane:

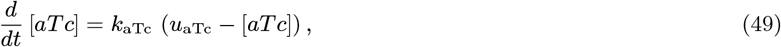

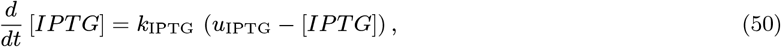

where the parameters *k*_aTc/IPTG_ are diffusion rates, whose values are reported in Table 1.

The deterministic model in (45)-(48) can be simplified by using a separation of time scales argument, as done in [32, 35]. Firstly, we can observe that equations (45) and (46) can be rewritten as

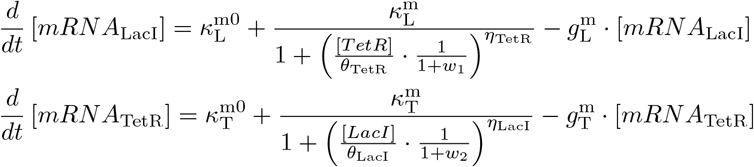

where 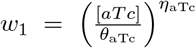 and 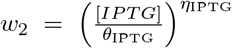. Now, taking into account that mRNA molecules are degraded faster than other molecules, we can obtain a quasi-steady state approximation by setting 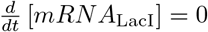 and 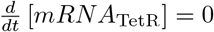, yielding

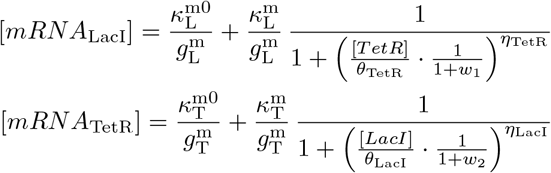

Assuming that LacI and TetR proteins degrade at the same rate, that is 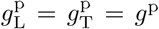, and using the dimensionless state variables 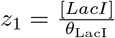 and 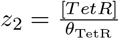, and dimensionless time *t′* = *g*^p^*t*, we obtain from (47):

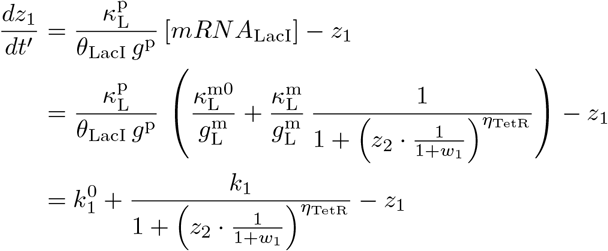

where

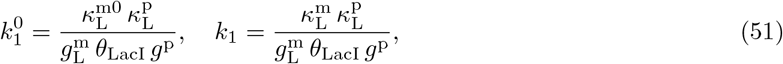

and from (48):

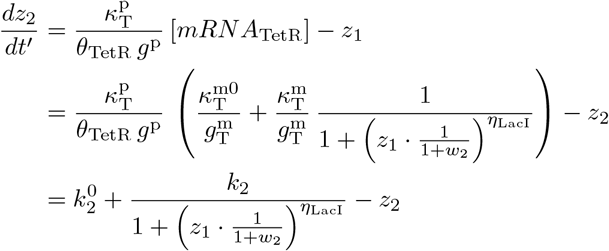

where

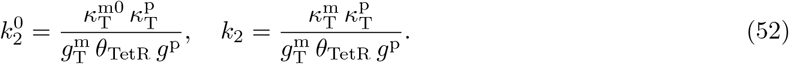

#### 7.8 Agent-based simulations in BSim

BSim is an agent-based simulator explicitly designed for the simulation of bacterial populations [22, 23]. In this environment the chemicals’ spatial distribution and the bio-mechanics of the cells are also considered. Specifically, it is possible to mimic a microfluidic platform, where cells grow and move, and where chemicals diffuse into the environment.

The BSim implementation presented in [23] has been extended here to include also stochastic simulations of the biochemical processes taking place inside the cells. Specifically, we implemented the SDE-based algorithm presented in [39]. Formally, for each cell we solved a system of equations of the form

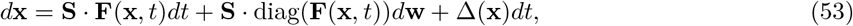

where **x** = [*mRNA*_LacI_], [*mRNA*_TetR_], [*LacI*], [*TetR*], [*aTc*], [*IPTG*]^┬^ is the state vector, **w** is a vector of independent standard Wiener processes, **S** = [**s**_1_, ..., **s**_6_] is the stoichiometry matrix and **F**(**x**) = [*a*_1_(**x**), *..., a*_6_(**x**)]^┬^, where *a*_*j*_(**x**) is the propensity function of the *j*-th pseudo-reaction defined as in STAR Methods 7.7. Notice that the term Δ(**x**)*dt* models the diffusion dynamics of chemicals across the cell membrane, assumed to be deterministic.

The microfluidic platform we considered consists of a microfluidic device, a microscopy, a computer and an actuation system. The microfluidic device contains a chamber where cells are trapped and grow. Specifically, we simulated the microfluidic chip described in [40], where the growth chambers’ dimensions are 40*μ*m × 30*μ*m × 1*μ*m. This allows bacteria to grow only in a mono-layer structure. The chamber is connected to two perfusion channels, which bring nutrients and inducers to the cellular population. The flow of the fluids sent to the chambers is governed by two syringes, each one containing a mixture of growth media and one inducer. By adjusting the relative height of the syringes, it is possible to change the amount of inducers delivered to the cells. The architecture is completed by a computer, controlling the syringes depending on the output of a control algorithm, and a microscopy, necessary to take images whose analysis results in the computation of the error.

The bio-mechanical parameters of the cells (i.e., cells’ growth rate) have been set as in [23].

Moreover, we have taken into account the following realistic constraints on the experimental platform [29] :

1. the state of the cells cannot be measured more often than 5 min, to avoid excessive photo-toxicity;
2. there is a time delay of 40 s on the actuation of the control inputs due to the time that the flow of the chemical inducers takes to reach the chambers in the microfluidic chip where cells are hosted;
3. the minimum time interval between two consecutive control inputs cannot be smaller than 15 min, to limit excessive osmotic stress on the cells;
4. the maximum duration of any experiment cannot exceed 24 hours (1440 min), to avoid substantial cell mutations during the experiments.

The specific implementation of the microfluidic device we adopted also introduces constraints on the possible classes of input signals *u*(*t*) = [*u*_aTc_(*t*), *u*_IPTG_(*t*)]^┬^ that can be generated by the actuators. We consider two possible implementations:

1. a T-junction, which limits *u*_aTc_ and *u*_IPTG_ to be mutually exclusive and with fixed amplitudes, that is, *u* is either equals to [*U*_aTc_, 0]^┬^, or to [0, *U*_IPTG_]^┬^;
2. a Dial-A-Wave (DAW) system [37], which constraints *u*_aTc_ and *u*_IPTG_ to be in a convex combination. Namely, given *u*_aTc_ ∈ [0, *U*_aTc_] we have

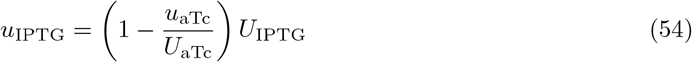

In the above equations, *U*_aTc_ and *U*_IPTG_ denote the concentrations of the inducers in the reservoirs, and we assume that they can be varied at the beginning of the experiments in the intervals [0, 100] and [0, 1], respectively. The upper bounds for *U*_aTc_ and *U*_IPTG_ are selected to avoid excessive stress on cells and are the same as those used *in vivo* in [27].

#### 7.9 Realistic in silico implementation of the controllers

To meet the specific dynamical features of the genetic toggle-switch implementing the required bistable memory mechanism in reversible differentiable cells (Section 2.4), we propose modified versions of the ideal relay controller and the PI controller proposed in Section 2.2.

Due to higher dimensional dynamics and unavoidable uncertainties affecting the system, it is not possible to define precise boundaries between the regions of attraction of the equilibrium points *A*_*i*_ and *B*_*i*_ for each cell. For this reason, we define conservative estimation of the regions of attraction common to all cells. Specifically, we defined 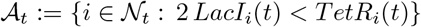 and 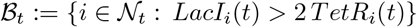, where *LacI*_*i*_ and *TetR*_*i*_ denote the concentrations of the molecules inside the *i*-th cell. Moreover, since in reality the union of these two sets does not cover the entire set 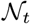 of cells in the consortium, cells not belonging to 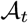 and 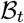 are in an *uncertain* state. We define this uncertain set of cells as 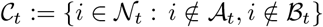 and denote with 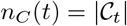 its cardinality at time *t*. For this reason, the error signals *eA* and *eB* are not complementary (i.e., *e*_*A*_(*t*) ≠ −*e*_*B*_(*t*)) and we need to consider both in the design of the control actions.

##### Relay controller

The relay controller consists of two mutually exclusive inputs with fixed amplitudes, which are applied to the system depending on the current value of the error signals *e*_*A*_(*t*) and *e*_*B*_(*t*). Specifically, at any time *t*, it is applied the input that causes max{|*e*_*A*_(*t*)|, |*e*_*B*_(*t*)|} to decrease. More formally, the control input *u*(*t*) = [*u*_aTc_(*t*), *u*_IPTG_(*t*)]^┬^ is chosen as

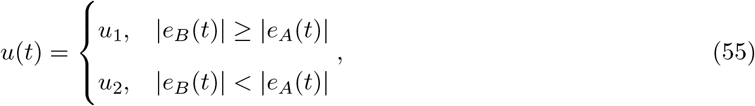

where

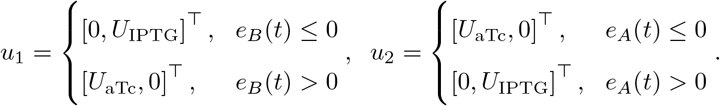

In this way, when *e*_*B*_ > 0 (*e*_*B*_ ≤ 0) a high value of aTc (IPTG) is applied, that, by promoting LacI (TetR), causes *e*_*B*_ to decrease (increase). Similar arguments hold for *e*_*A*_.

The control parameters *U*_aTc_ and *U*_IPTG_ were tuned empirically by trial-and-error. The numerical values used in each simulation are reported in the caption of the relative figure.

##### PI controller

The PI controller is implemented using a Dial-A-Wave system [37]. It consists of two control actions, each acting to decrease either of the control errors. Formally, we have:

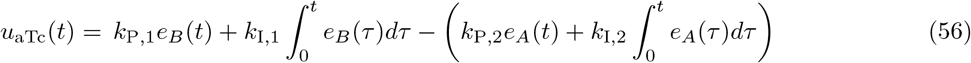

where *k*_P,j_ and *k*_I,j_, *j* = 1, 2, are the proportional and integral control gains. Note that, due to the constraints on the actuation system, *u*_IPTG_ is defined by (54).

The proportional and the integral gains, and the maximum concentrations of the inducers *U*_*aTc*_ and *U*_*IPTG*_ are empirically tuned by trial-and-error. The numerical values used in each simulation are reported in the caption of the relative figure.

Moreover, to improve performance, the control algorithm is complemented with an anti-windup scheme (that sets to zero the internal state of the integrator whenever the error signals are equal to 0 or change sign) and a dynamic saturation defined as:

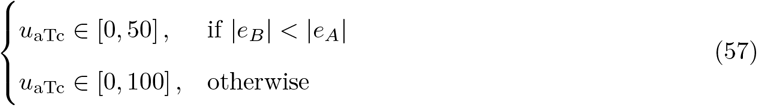

Similarly for *u*_IPTG_ as defined by (54).

#### 7.10 Noise magnitude due to flush-out effects

The fluctuations in the steady-state response of the closed-loop system (see for example Figure 6) are mostly due to cells flowing out of the microfluidic chamber as they grow. The magnitude *ε* of this noise is indeed proportional to the ratio between the flow rate *Q* of cells exiting the chamber and the total number *N* of cells therein. That is, *ε* ∝ *Q/N*, where *Q* = *v* · *A*; being *A* the cross-section area of the chamber apertures and *v* the flow velocity, that depends on the cell growth rate. Without loss of generality, assume that, as is usual in microfluidic applications, cells in the chamber are organized in a single-layer and that, on average, the area occupied by a cell can be approximated by a circle of radius *r*. Moreover, let us assume that the chamber has a square geometry with side *ℓ* with two open sides, as in Figure 5. Then, the flow rate *Q* can be approximated as 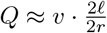. Moreover, the total number of cells in the chamber is *N* = *ℓ*^2^/(*πr*^2^). Therefore, in conclusion, we have that

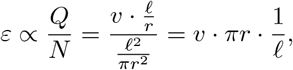

that is, the magnitude of the noise is inversely proportional to the size of the culture chamber. We validated the formula numerically by comparing the noise levels between chambers of different sizes (compare Figure 6 and Supplemental Figure 11). Specifically, we observed that, as expected, the average variation of the ratios at steady state in the chamber 40*μm* × 50*μm* wide (0.009) was three times lower than the one in a chamber where both the dimensions were scaled down to a third (0.03) (see also Supplemental Video 2).

Note that when the cells under observation flow out of the field of view from all the four sides of the chamber (a common situation when higher magnification factors are used on the microscope), the same result still holds but multiplied by a factor of 2.

## Supplemental Videos

- Supplemental Video 1: Animations of the *in silico* control experiments in BSim reported in Figure 6.
- Supplemental Video 2: Animations of the *in silico* control experiments in BSim reported in Supplemental Figure 11, with a smaller growth chamber.

## Data and code availability

The code used for all simulations is available at https://github.com/diBernardoGroup/Ratiometric-Control-of-bacterial-populations

## 8 Supporting Information

**Figure 8:**
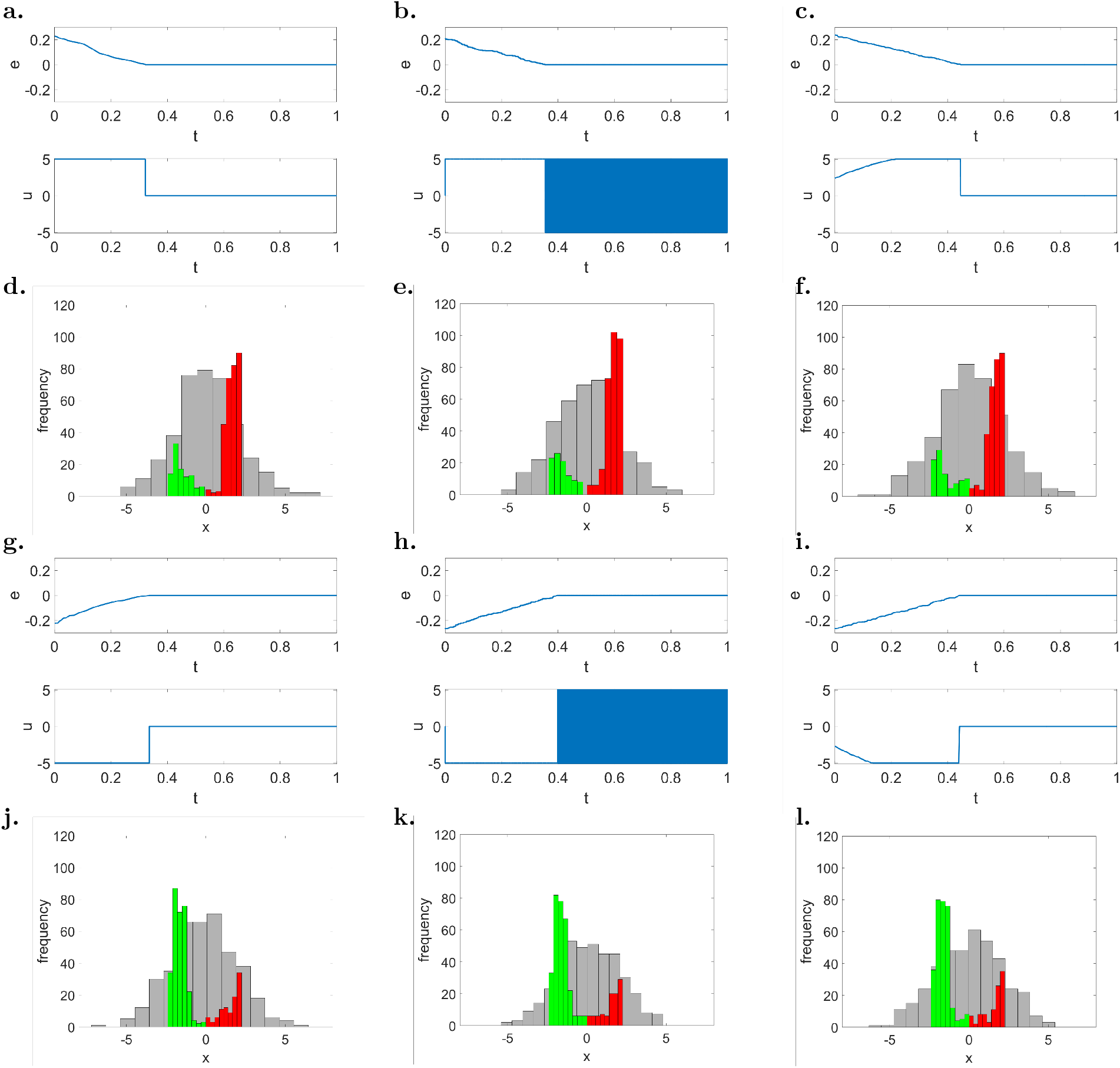
Feedback control strategies are effective to balance two *asymmetric* groups of controllable reversible cells to 1:3 ratio (*r*_d_ = 0.75) and 3:1 ratio (*r*_d_ = 0.25). (**a-c, g-i**) Evolution of the error signal *e*(*t*) and of the control input *u*(*t*) for (**a, g**) the first implementation of the relay controller (22), (**b, h**) the second implementation of the relay controller (24), (**c, i**) the PI controller (26)-(27). (**d-f, j-l**) Distribution of the cells state at the beginning of the simulation (*t* = 0 a.u., gray histogram) and at steady state (*t* = 1.0 a.u., green and red histograms), for (**d, j**) the first implementation of the relay controller, (**e, k**) the second implementation of the relay controller, (**f, l**) the PI controller. Desired ratios are set to (**a-f**) *r*_d_ = 0.75 (1:3 ratio) and (**g-l**) *r*_d_ = 0.25 (3:1 ratio). The green and red bars in panels **d-f** and **j-l** correspond to cells in the basin of attraction of *A*_*i*_ and *B*_*i*_, respectively. The maximum control input is set to *ū* = 5 and the gains of the PI controller has been set to *k*_P_ = 30 and *k*_I_ = 10. All cells (*N* = 400) have initial conditions *x*_*i*_(0) drawn from the normal random distribution 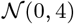, and the parameters *η*_*i*_ are drawn with uniform distribution from the interval [1, 5], therefore all cells are controllable, as no monostable 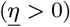 and no unswitchable 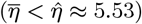 cells are present in the population.

**Figure 9:**
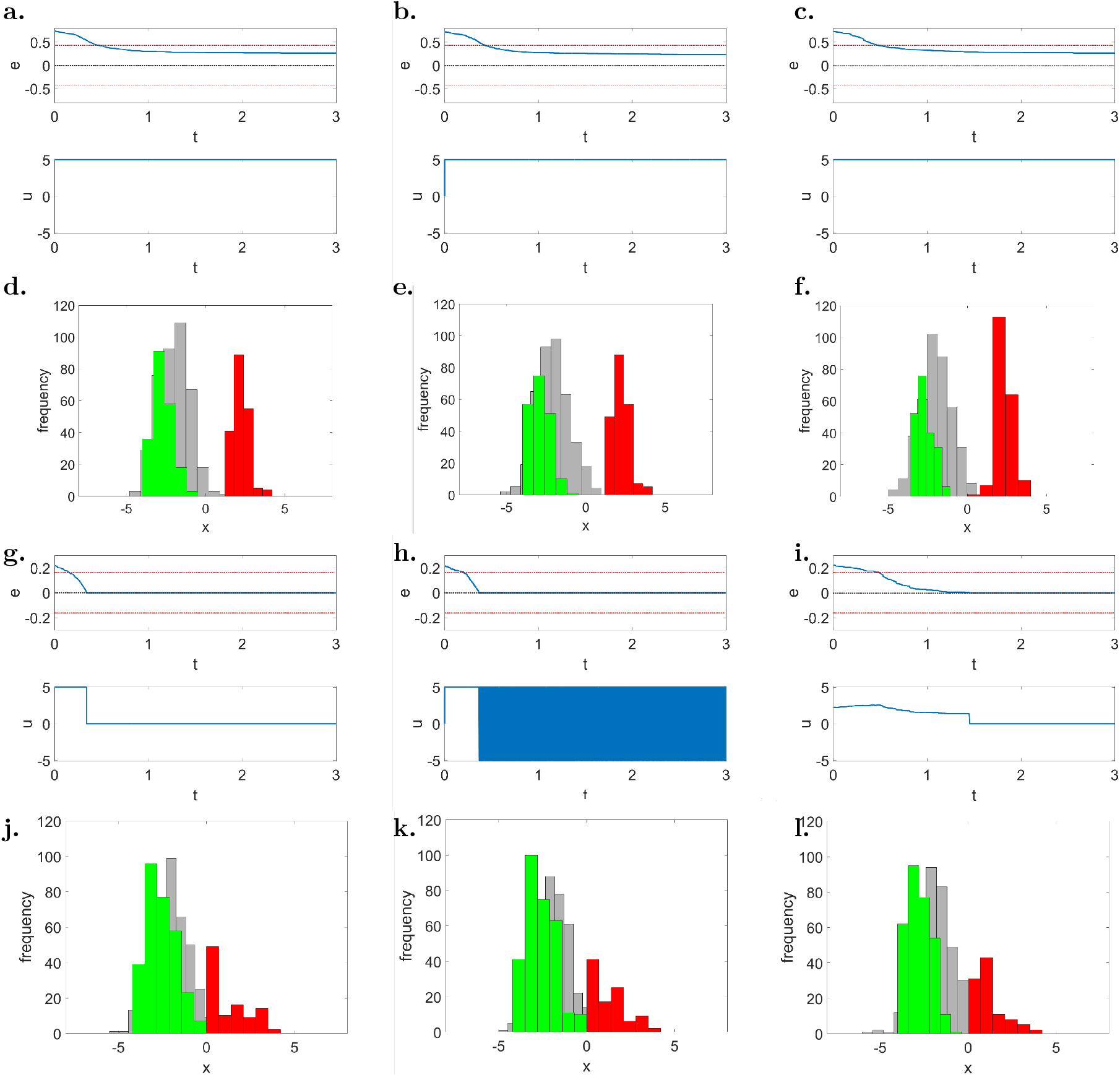
Balance to *asymmetric* ratios is achieved with a residual steady-state error in the presence of uncontrollable cells. (**a-c, g-i**) Evolution of the error signal *e*(*t*) and of the control input *u*(*t*) for (**a, g**) the first implementation of the relay controller (22), (**b, h**) the second implementation of the relay controller (24), (**c, i**) the PI controller (26)-(27). (**d-f, j-l**) Distribution of the cells state at the beginning of the simulation (*t* = 0 a.u., gray histogram) and at steady state (*t* = 3.0 a.u., green and red histograms), for (**d, j**) the first implementation of the relay controller, (**e, k**) the second implementation of the relay controller, (**f, l**) the PI controller. Desired ratios are set to (**a-f**) *r*_d_ = 0.75 (1:3 ratio) and (**g-l**) *r*_d_ = 0.25 (3:1 ratio). The green and red bars in panels **d-f** and **j-l** correspond to cells in the basin of attraction of *A*_*i*_ and *B*_*i*_, respectively. The maximum control input is set to *ū* = 5 and the gains of the PI controller are set to *k*_P_ = 30 and *k*_I_ = 10. All cells (*N* = 400) have initial conditions *x*_*i*_(0) drawn from the normal random distribution 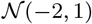, and the parameters *η*_*i*_ are drawn with uniform distribution from the interval [−1, 14], therefore both monostable 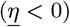 and unswitchable 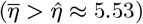 cells can be present in the population. The steady-state errors observed in the *in silico* experiment are equal to **(a)** 0.265, **(b)** 0.235, **(c)** 0.265, and **(g-i)** 0. Note that all the observed errors are below the theoretical upper bound on the control error estimated using (5) as 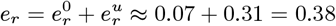. (depicted in the panels **a-c** and **g-i** as red dashed lines).

**Figure 10:**
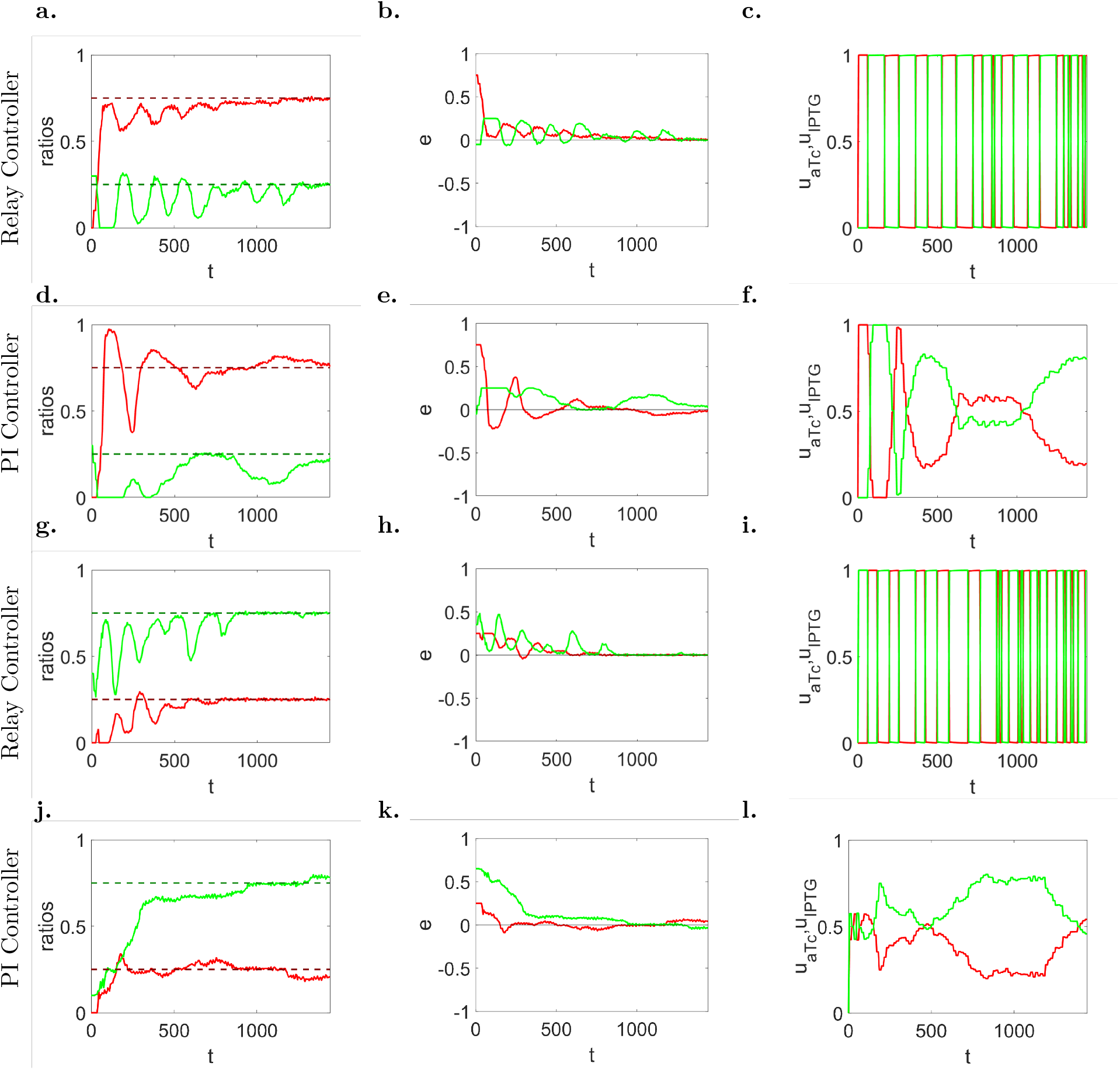
Cooperative production of two monomers to *asymmetric* population ratios can be achieved by means of feedback ratiometric controllers in microfluidics. (**a, d, g, j**) Evolution in time of populations’ ratio *r*_*A*_ (solid green line) and *r*_*B*_ (solid green line) with their respective desired reference values in dashed lines, (**b, e, h, k**) of the error signals *eA* (solid green line) and *eB* (solid red line), and (**c, f, i, l**) inducer control signals *u*_aTc_ (solid red line) and *u*_IPTG_ (solid green line), normalized to their maximum values *U*_aTc_ and *U*_IPTG_, respectively. Desired ratios are set to (**a-f**) *r*_d_ = 0.75 (1:3 ratio) and (**g-l**) *r*_d_ = 0.25 (3:1 ratio). (**a-c, g-i**) Parameters of the relay control (55): *U*_aTc_ = 60 ng/mL, *U*_IPTG_ = 0.5 mM. (**d-f, j-l**) Parameters of the PI controller (56): *U*_aTc_ = 60 ng/mL, *U*_IPTG_ = 0.5 mM, *k*_P,1_ = 60, *k*_P,2_ = 0.75, *k*_I,1_ = 1.5, *k*_I,2_ = 0.05. Cells (about 200) in the simulated microfluidic chamber (with dimensions 40 *μ*m × 50 *μ*m × 1 *μ*m) have all the same parameters’ value, and their evolution has been obtained using the agent-based simulator BSim [22, 23].

**Figure 11:**
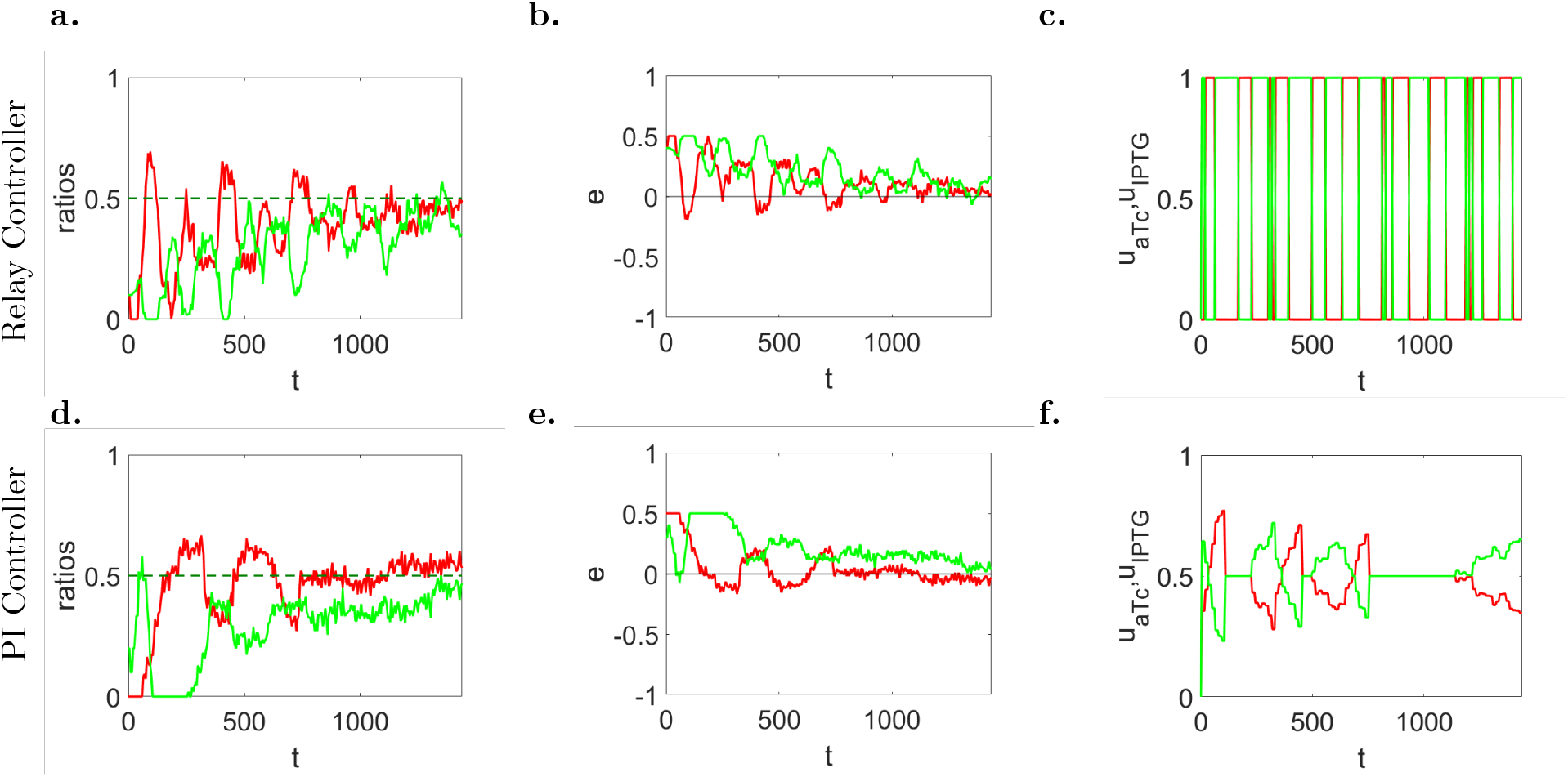
Cooperative production of two monomers to 1:1 population ratio can still be achieved in *smaller* growth chambers. Larger noise fluctuations due to cells flowing out of the microfluidic chambers are expected in this situation. (**a,d**) Evolution in time of populations’ ratio *r*_*A*_ (solid green line) and *r*_*B*_ (solid green line) with their respective desired reference values in dashed lines (*r*_d_ = 0.5), (**b,e**) of the error signals *e*_*A*_ (solid green line) and *e*_*B*_ (solid red line), and (**c,f**) inducer control signals *u*_aTc_ (solid red line) and *u*_IPTG_ (solid green line), normalized to their maximum values *U*_aTc_ and *U*_IPTG_, respectively. (**a-c**) Parameters of the relay control (55): *U*_aTc_ = 60 ng/mL, *U*_IPTG_ = 0.5 mM. (**d-f**) Parameters of the PI controller (56): *U*_aTc_ = 100 ng/mL, *U*_IPTG_ = 1 mM, *k*_P,1_ = 100, *k*_P,2_ = 1.5, *k*_I,1_ = 1.5, *k*_I,2_ = 0.05. The simulated microfluidic chamber (with dimensions 13.3 *μ*m× 16.7 *μ*m× 1 *μ*m) can host a cell population of about 30 cells. All simulations were obtained using the agent-based simulator BSim [22, 23]. (See also Supplemental Video 2.)

**Figure 12:**
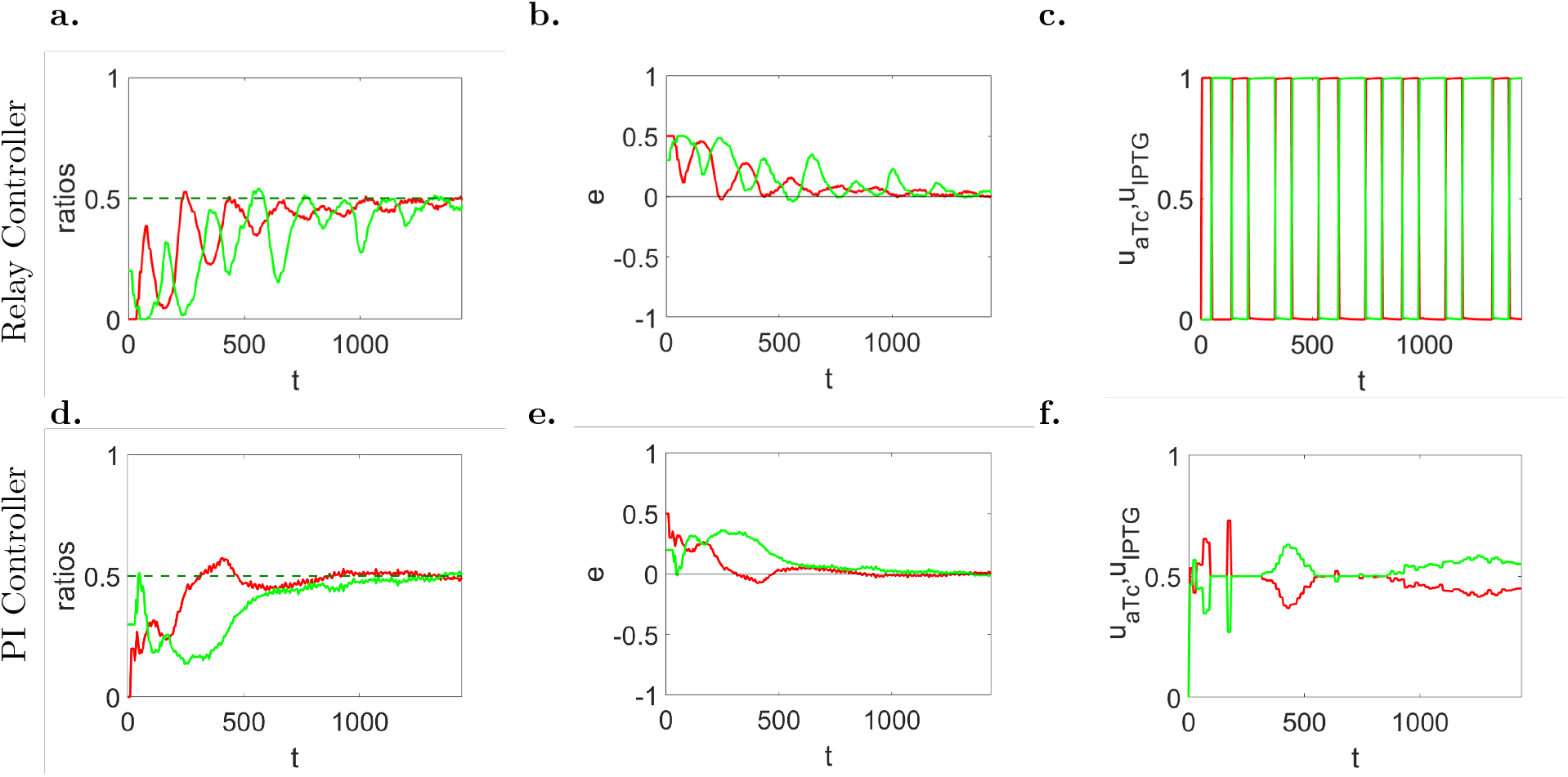
Cooperative production of two monomers to 1:1 population ratio can be achieved in a *heterogeneous* cell population by means of feedback ratiometric controllers in microfluidics. (**a,d**) Evolution in time of populations’ ratio *r*_*A*_ (solid green line) and *r*_*B*_ (solid green line) with their respective desired reference values in dashed lines (*r*_d_ = 0.5), (**b,e**) of the error signals *e*_*A*_ (solid green line) and *e*_*B*_ (solid red line), and (**c,f**) inducer control signals *u*_aTc_ (solid red line) and *u*_IPTG_ (solid green line), normalized to their maximum values *U*_aTc_ and *U*_IPTG_, respectively. (**a-c**) Parameters of the relay control (55): *U*_aTc_ = 60 ng/mL, *U*_IPTG_ = 0.5 mM. (**d-f**) Parameters of the PI controller (56): *U*_aTc_ = 100 ng/mL, *U*_IPTG_ = 1 mM, *k*_P,1_ = 100, *k*_P,2_ = 1.5, *k*_I,1_ = 1.5, *k*_I,2_ = 0.05. The simulated microfluidic chamber (with dimensions 40 *μ*m × 50 *μ*m × 1 *μ*m) can host a cell population of about 200 cells. Each time a cell divides, the parameters of the daughter are drawn from a normal distribution centered in the nominal value, and with variance equal to its 20%. All simulations were obtained using the agent-based simulator BSim [22, 23].

## Notes

### Competing Interest Statement

The authors have declared no competing interest.

